# High-throughput RNA isoform sequencing using programmable cDNA concatenation

**DOI:** 10.1101/2021.10.01.462818

**Authors:** Aziz M. Al’Khafaji, Jonathan T. Smith, Kiran V Garimella, Mehrtash Babadi, Moshe Sade-Feldman, Michael Gatzen, Siranush Sarkizova, Marc A. Schwartz, Victoria Popic, Emily M. Blaum, Allyson Day, Maura Costello, Tera Bowers, Stacey Gabriel, Eric Banks, Anthony A. Philippakis, Genevieve M. Boland, Paul C. Blainey, Nir Hacohen

**Author notes:** Corresponding authors, Aziz M. Al’Khafaji, Kiran V Garimella, Mehrtash Babadi, Paul C. Blainey Nir Hacohen. These authors contributed equally (Aziz M. Al’Khafaji, Jonathan T. Smith, Kiran V Garimella, Mehrtash Babadi).

## Abstract

Alternative splicing is a core biological process that enables profound and essential diversification of gene function. Short-read RNA sequencing approaches fail to resolve RNA isoforms and therefore primarily enable gene expression measurements - an isoform unaware representation of the transcriptome. Conversely, full-length RNA sequencing using long-read technologies are able to capture complete transcript isoforms, but their utility is deeply constrained due to throughput limitations. Here, we introduce MAS-ISO-seq, a technique for programmably concatenating cDNAs into single molecules optimal for long-read sequencing, boosting the throughput >15 fold to nearly 40 million cDNA reads per run on the Sequel IIe sequencer. We validated unambiguous isoform assignment with MAS-ISO-seq using a synthetic RNA isoform library and applied this approach to single-cell RNA sequencing of tumor-infiltrating T cells. Results demonstrated a >30 fold boosted discovery of differentially spliced genes and robust cell clustering, as well as canonical PTPRC splicing patterns across T cell subpopulations and the concerted expression of the associated hnRNPLL splicing factor. Methods such as MAS-ISO-seq will drive discovery of novel isoforms and the transition from gene expression to transcript isoform expression analyses.

## Main

While RNA sequencing has dramatically accelerated our understanding of biology, accurate quantification and discovery of RNA isoforms, especially at single-cell resolution, remains a steep challenge^1^. Alternative splicing is a core regulatory process that modulates the structure, expression, and localization of proteins through differential exon and/or UTR splicing during transcript maturation in eukaryotes. Beyond being an integral component of cellular/organismal development and homeostatic maintenance, alternative splicing is implicated in a wide range of pathologies with hallmark isoforms being linked to cardiovascular, neurological, and immunological diseases^2,3^. Additionally, mutated and/or dysregulated splicing factors make up a major class of phenotypic alterations associated with tumor progression and therapeutic resistance^4^.

High-throughput full-length RNA isoform identification and quantification remain elusive goals for single-cell and bulk studies as the necessary read lengths (>5 kb) and depths (>2×10^7^ reads) are not easily attainable by existing sequencing platforms (Supplementary Fig. 1). For example, short-read sequencing platforms (e.g. Illumina) achieve more than sufficient throughput (>1×10^9^ reads) but are hindered by limited read lengths (50-600 bp) which are inadequate to span the majority of human transcripts (∼ 1.6 ± 1.1 kb, Supplementary Fig. 2). As a result, individual short reads often fail to span successive splice sites, impairing efforts to correctly identify alternative transcript isoforms^5^. A recently developed short-read sequencing approach, Smart-seq3, enhances isoform detection by enabling single-molecule reconstruction via integration of reads from products with the same 5’ unique molecular identifier (UMI)^6^. However, due to the 5’ coverage bias of Smart-seq3, the vast majority of transcript molecules are only partially reconstructed, resulting in poor isoform identification and discovery. Conversely, the long-read platforms Oxford Nanopore (ONT) and Pacific Biosciences (PacBio) enable full-length RNA isoform sequencing but suffer from comparatively low read throughput at high costs. Early limitations in raw base calling accuracy on long-read platforms (error rates of 10-15%) have been significantly mitigated by circularized consensus sequencing (CCS, also referred to as HiFi) and consensus generation strategies for individual library molecules^7,8^. On the PacBio Sequel IIe platform, consensus base quality reaches Phred-scale quality of ∼Q30 around 10 circular passes, with subsequent consensus reads providing marginal further gains. For the current Sequel IIe instrument and SMRT Cell 8M chemistry, the optimal library size for reaching ∼10 circular passes is 15-20 kb. As the length register of transcripts is on average substantially shorter (100 bp-5 kb), CCS of individually circularized cDNA molecules yields an excessive number of circular passes (50-60), effectively wasting sequencing potential (Supplemental Fig. 3).

To maximize the sequencing throughput on the PacBio platform, we developed a method for the programmable concatenation of DNA fragments into long composite sequence library molecules, **M**ultiplexed **A**rray**s S**equencing (MAS-seq, Fig. 1a). When MAS-seq is used for sequencing transcript isoforms, we term the approach MAS-ISO-seq. The protocol begins by depleting TSO priming artifacts via streptavidin/biotin selection. Purified cDNAs then undergo parallel PCRs, which serve to both increase cDNA yield and append reaction specific deoxy-uracil (dU) containing barcode adapters. Through the use of dU digestion followed by barcode-directed ligation of cDNAs, MAS-ISO-seq generates long concatenated cDNA arrays with a narrow length distribution that allows for both accurate consensus sequencing and more optimal capacity utilization of the PacBio long-read platform. To drive accurate and specific hybridization, the ligation barcode adapters are designed to be 15 bp in length with each having a hamming distance of 11 from all other barcodes^9^. In combination with upstream depletion of TSO priming artifacts via streptavidin/biotin selection, MAS-ISO-seq boosts the sequencing throughput to approximately 40 million full-length transcripts per SMRT Cell 8M flow cell, a >15-fold increase over CCS corrected read counts (Fig. 1b).

**Fig. 1:**
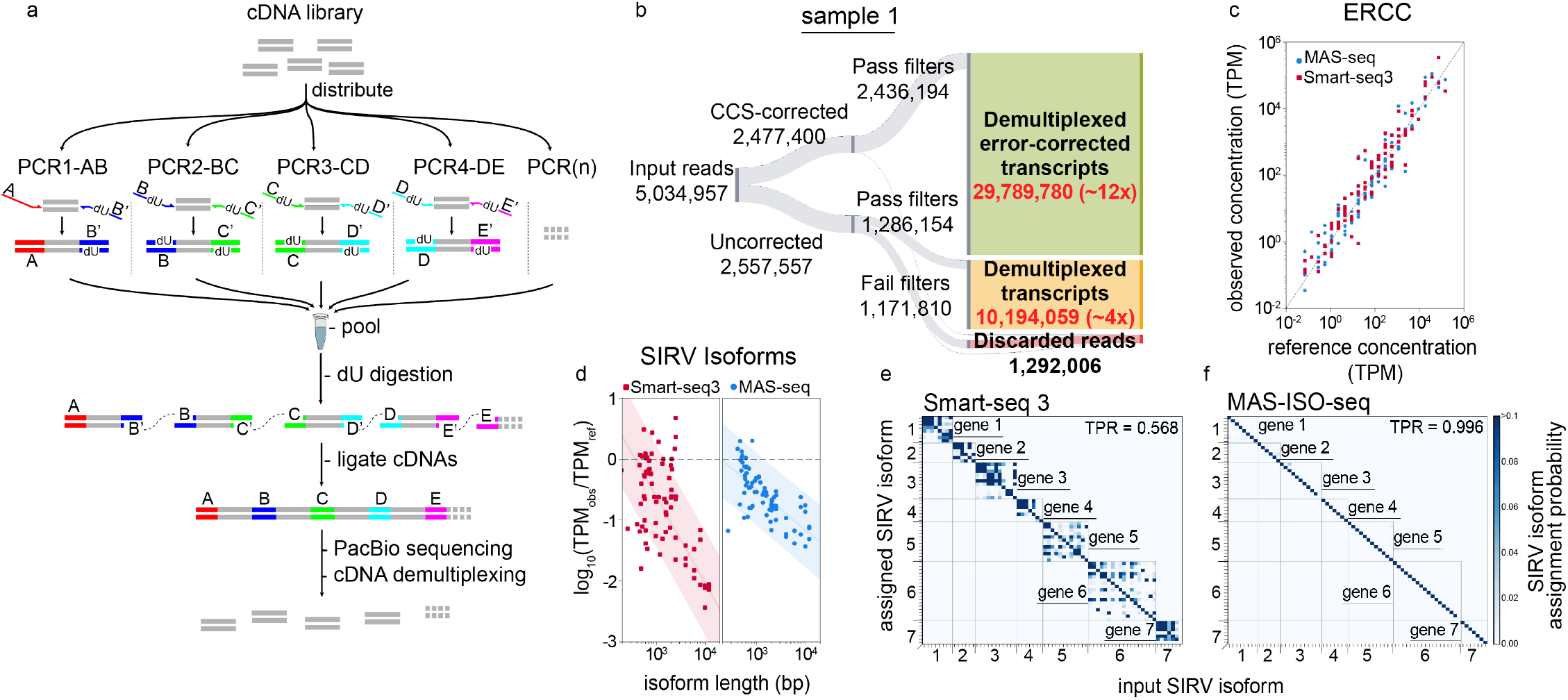
MAS-ISO-seq workflow and experimental validation using synthetic RNA isoforms. **(a)** Schematic of the MASISO-seq intramolecular cDNA multiplexing workflow. **(b)** Sankey diagram reporting MAS-ISO-seq run yield of sample 1 at various stages of processing. **(c)** Observed ERCC concentrations as measured in MAS-ISO-seq and Smart-seq3 experiments vs. reference concentrations (R-squared > 0.95 for both). **(d)** Log-ratio of observed to reference concentrations of short and long SIRV isoforms in SIRV-Set 4 vs. transcript length for Smart-seq3 and MAS-ISO-seq. **(e)** Isoform identification confusion matrix for SIRV isoforms as measured by Smart-seq3 reconstructions and **(f)** MAS-ISO-seq reads.

To demonstrate the throughput gain and utility of MAS-ISO-seq, we performed a 15-member cDNA ligation from two 5’ single-cell gene expression cDNA libraries (10x Genomics) of tumor-infiltrating CD8+ T cells. As expected, we observed a narrowly distributed ∼15-fold increase in cDNA library size after ligation (Supplementary Fig. 4). MAS-ISO-seq libraries underwent standard CCS library preparation and were sequenced on the PacBio Sequel IIe. Sequenced libraries exhibited corrected read length and circular pass count distributions more comparable to whole-genome CCS data than the standard isoform sequencing method, Iso-Seq, as expected due to longer concatenated library lengths (Supplementary Fig. 5).

The sequential pattern of distinct MAS-ISO-seq adapters provides landmarks for effective cDNA segmentation as well as constraints for detecting malformed or otherwise defective sequences. MAS-ISO-seq adapters also enable utilization of CCS uncorrected reads which are otherwise discarded. To exploit these signals, we developed a composite profile hidden Markov model, *Longbow*, for the probabilistic annotation and optimal segmentation of each MAS-ISO-seq read via maximum *a posteriori* state path. *Longbow* is robust to the presence of a high per-base error rate (Methods). Across both single-cell MAS-ISO-seq libraries, 98.9-99.0% of CCS corrected reads and 43.0-50.3% of CCS uncorrected reads were found to segment consistently. To maximize precision, segmentation results inconsistent with our expected array structure (i.e. off-subdiagonal elements of the matrices in Supplementary Fig. 6a,b) were filtered out (Supplementary Fig. 6c,d). A plurality of filtered reads (Sample 1: 31.14%, Sample 2: 38.18%) were found to contain fully-formed 15-element arrays. Arrays with fewer than 15 cDNAs were more prevalent in uncorrected reads than corrected (Supplementary Fig. 7,8). Nonetheless, the vast majority of demultiplexed reads from these partial arrays post-filtering still contained consecutive adapter sequences, a poly(A) tail, and had a high mapping quality to the genome (95.53%, Supplementary Fig. 7,8). After final filtering and segmentation across both samples, we obtained ∼22-29M CCS-corrected transcripts (a ∼12-14 fold yield increase over the number of CCS corrected reads) and ∼10-13M CCS-uncorrected transcripts (an additional ∼4-8 fold increase) for a total 16-22 fold increase as compared to standard CCS corrected reads (Fig. 1b, Supplementary Fig. 9).

To validate the ability of MAS-ISO-seq to faithfully capture full-length RNA isoforms, we performed full-length RNA sequencing of the Lexogen SIRV-Set 4, a synthetic mixture of Spike-In RNA Variants (SIRVs) containing 69 RNA isoforms of varying lengths and equal molarity across 7 “genes”, 15 long 4-12kb SIRVs, and 92 ERCC RNA standards with concentration spanning 6 orders of magnitude^10^. Smart-seq3 sequencing of the SIRV-Set 4 library was performed in parallel to compare short-read isoform reconstructions to our high-throughput long-read sequencing approach. While quantification of ERCC standards was broadly similar overall between both protocols (Fig. 1c), long isoforms showed markedly reduced length bias in MAS-ISO-seq and Iso-Seq vs. Smart-seq3 (Fig. 1d, Supplementary Fig. 10). Smart-seq3 isoform reconstructions exhibited substantial ambiguity in assigning reconstructed transcripts to a specific known isoform (∼43% error rate) (Fig. 1e). In contrast, MAS-ISO-seq allows direct identification of transcript isoforms without the need for *in silico* reconstruction, and hence leads to virtually unambiguous isoform assignment (∼0.4% error rate, Fig. 1f).

To characterize the performance of MAS-ISO-seq for single-cell RNA sequencing, we performed 10x Genomics 5’ single-cell gene expression on tumor-infiltrating CD8+ T cells. Using the standard 5’ single-cell gene expression protocol, we generated both standard short-read and MAS-ISO-seq long-read libraries from the same full-length cDNA library. After conventional QC filtering steps and removal of doublets and primary tumor cells (Methods), we obtained 6,260 CD8+ T cells containing median 3,211 UMIs/cell (short read data) and 1,538 UMIs/cell (long read data). Sequencing saturation was higher for the short-read run, 3.71 reads/UMI (short) vs. 1.37 reads/UMI (long). Despite large discrepancies in sequencing depth between short and long-read approaches and quantification methodologies (Methods), cell clustering and gene expression were highly concordant (Fig. 2a, adjusted Rand index = 0.78; Fig. 2b, concordant gene count saturation curves; Supplementary Fig. 11, R-squared = 0.85). A common set of T cell transcriptional states ranging from stem cell-like to terminally differentiated were observed in both datasets.

**Fig. 2:**
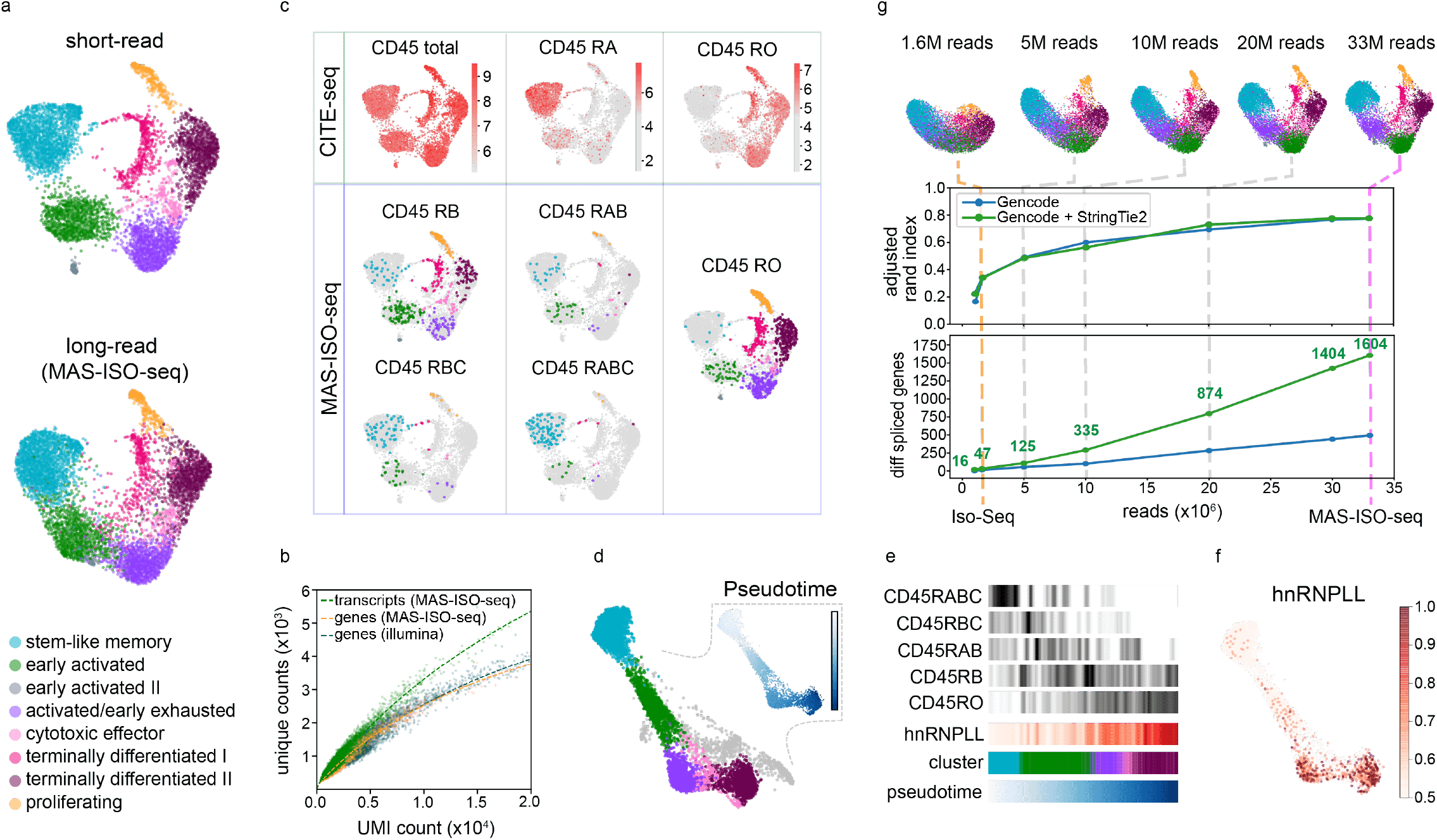
Single-cell isoform-resolved sequencing of primary human CD8+ T cells with MAS-ISO-seq. **(a)** UMAP embedding of single-cell gene expression of 6,260 CD8+ T cells from short or long-read analyses; the long-read UMAP is annotated with the cell identities determined from the short-read data. **(b)** Scatter plot of unique gene or transcript counts in cells vs. UMI counts for short-read (Illumina) and long-read (MAS-ISO-seq). **(c)** CD45 (*PTPRC*) isoform analysis using either CITE-seq or MAS-ISO-seq. **(d)** Force directed graph of CD8+ T cells with insets depicting pseudotime progression and differential CD45 isoform expression along the pseudotime axis. **(e)** Levels of isoforms along pseudotime and in each cluster. **(f)** Expression of hnRNPLL along the pseudotime progression; **(g)** Downsampling analysis of MAS-ISO-seq reads; (top) evolution of UMAP embedding vs. depth; (middle) adjusted Rand index (ARI) between short-reads reference annotations and downsampled long reads vs. depth; (bottom) number of statistically significant differentially spliced genes vs. depth.

Leveraging the distinct splicing patterns of CD45 (*PTPRC*) over the course of T cell differentiation, we performed orthogonal validations of CD45 isoform expression at the protein level using CITE-seq and compared them to the mRNA levels measured with MAS-ISO-seq^11^. CD45 isoform expression between these two modalities was highly concordant (Fig. 2c). Notably, mRNA measurements were more granular in their ability to resolve the multiple CD45 isoforms present (RO, RA, RAB, RB, RBC) as compared to the antibody-based CITE-seq approach. This is due to the single epitope specificity of antibodies which do not enable discrimination of closely related isoforms^12^. For example, the CD45 RA antibody cannot distinguish between CD45 RA and RAB. Pseudotime analysis revealed a continuum of T cell states leading from stem cell-like to activated to terminally differentiated. Canonical CD45 isoform expression and its associated splicing factor, hnRNPLL^11^, tracked clearly along this differentiation trajectory (Fig. 2d-f).

To quantify the impact of the sequencing depth gained by MAS-ISO-seq on cell typing and identification of differential spliced genes, we performed an *in silico* downsampling analysis from a single MAS-ISO-seq run. We processed each dataset identically using the same pipeline and computed the adjusted Rand index (ARI) between the cell clustering of the subsampled long-read dataset and the full short-read dataset as reference. We also determined the number of differentially spliced genes across the T cell subtypes for each dataset (Methods). Compared to the read depth expected from an Iso-Seq run (1.6M full-length transcripts), the throughput gain afforded by MAS-ISO-seq translates to 44% increase and saturation of ARI between short-read and long-read single-cell clustering and remarkably, 34-fold gain in identifying differentially spliced genes (multiple hypothesis testing correction with FDR < 0.05) (Fig. 2g; cluster-resolved results given in Supplementary Fig. 12). Notably, a plurality of the differentially spliced (DS) genes were distinct from the set of differentially expressed (DE) genes (Supplementary Fig. 13).

In summary, we detail and validate MAS-ISO-seq, a cDNA concatemerization method that boosts throughput of the PacBio long-read sequencing platform >15-fold to approximately 40 million reads per run. Using synthetic RNA isoforms as a ground truth library, we demonstrate that MAS-ISO-seq is far superior in confidently identifying RNA isoforms as compared to short-read approaches. Further, we leveraged MAS-ISO-seq to perform single-cell RNA isoform sequencing on human tumor-infiltrating CD8+ T cells. We validated our ability to accurately identify isoforms by resolving canonical CD45 isoform expression differences across the range of observed cell states and orthogonal protein isoform-based measurements. Through downsampling analyses, we demonstrate that the additional throughput afforded by MAS-ISO-seq enables robust cell clustering into known T cell differentiation states and substantially boosts the identification of differentially spliced genes. A related approach, HIT-scISOseq, leverages palindromic adapter sequences to drive ligation of an indeterminate number of cDNAs, enabling approximately 10 million transcript reads^13^. While producing four-fold lower yield as compared to MAS-ISO-seq, HIT-scISOseq additionally lacks the sequential array structure that MAS-ISO-seq exploits for accurate segmentation and identification of malformed arrays. Other concatenation approaches for targeted DNA sequencing exist which use Gibson Assembly or Golden Gate Assembly for array formation. These methods also demonstrate considerably lower throughput and lack the error robustness of MAS-seq arrays^14,15^.

Challenges impacting the RNA isoform sequencing field as a whole include cDNA synthesis artifacts, incomplete transcriptome references, and inaccurate transcriptome assembly software. We believe that the read throughput afforded by approaches such as MAS-ISO-seq will lower barriers to data generation and facilitate solving these related problems. The compatibility of MAS-ISO-seq with archived single-cell cDNA libraries from short-read platforms extensively applied in cell atlasing studies is poised to immediately facilitate isoform discovery and reference generation with cell type annotations at scale. Furthermore, MAS-ISO-seq will augment efforts related to fusion identification, neoantigen discovery, and TCR/BCR repertoire sequencing. Given the modular and scalable nature of the workflow, MAS-ISO-seq is positioned to co-evolve with long-read sequencing platforms, enabling even greater throughput as read lengths, yield, and per-base accuracy increase.

## Methods

### Patients consent and sample collection

Patients CD8^+^ T cells analyzed in this study were collected under the Dana-Farber/Harvard Cancer Center Institutional Review Board (DF/HCC protocol 11-181), and provided written informed consent prior to tissue collection.

### Single-cell and SIRV cDNA library preparation

#### Sample dissociation and FACS Sorting of CD3^+^CD8^+^ T cells

Using the human tumor dissociation kit (Miltenyi Biotec; Cat# 130-095-929), freshly isolated tumors were digested to obtain a single cell suspension. Tissue was placed into a 1.5mL Eppendorf tube containing 420μL of DMEM with 10% FCS, 42μL of enzyme H, 21μL of enzyme R, and 5μL enzyme A (provided with the kit). The tissue was minced using surgical scissors, and an additional 512μL of DMEM with 10% FCS was added to the tube (total volume of 1ml). Next the tissue was incubated for 15 min at 37°C, 350 rpm in a thermomixer (Eppendorf; F1.5). After incubation, the tissue was further digested using a 1 ml syringe plunger over a 50μm filter (Sysmex; Cat# 04-004-2327), making sure to wash the filter with media. Using ACK buffer (Gibco; Cat# A1049201), RBC lysis was performed and the sample was finally resuspended in DMEM with 10% FCS in order to count and determine the viability of the cells using a manual hemocytometer (Bright-line; Cat# 1492). Cells were then washed twice with cold PBSx1 and the cells were incubated with live/dead Zombie Violet Dye (Biolegend, 423114) for 15 min at RT as suggested by the manufacturer. The cells were then washed and resuspended with 1X PBS containing 1.5% FCS for cell surface labelling using a standard protocol for a 30 min at 4°C. A antibody panel was used to identify and sort the CD3^+^CD8^+^ T cell population: Human TrueStain FcX (Biolegend; Cat# 422302), PE anti-human CD45 (Biolegend; Cat# 304008), FITC anti-human CD3 (Biolegend; Cat# 317306), APC/Cyanine7 anti-human CD235a (Biolegend; Cat# 349116), and APC anti-human CD8a (Biolegend; Cat# 300912). Sorting of single live CD3^+^CD8+ T cells (gating on Zombie^low^, hCD235a^-^, hCD45^+^, hCD3^+^, hCD8^+^) was performed using a Sony MA900 cell sorter. Cells were sorted into a 15mL tube containing DMEM with 10% FCS. After sorting, tubes with sorted cells were vortexed briefly, spun down at 1500rpm, 4°C for 5 minutes, resuspended, and counted for yield.

### TotalSeq-C staining and Single-cell RNA sequencing procedure

Sorted CD3^+^CD8^+^ T cells were washed and resuspended with staining buffer (PBSx1 + FCS 2.5% + 2mM EDTA). Next TruStain FcX (FC blocker, Biolegend; Cat# 422301) was added and the sample was incubated for 10 min at 4°C. After incubation with FcX blocker, the cells were washed with staining buffer once and spun down at 1500rpm, 4°C for 5 minutes. The cells were then incubated for 20 min at 4°C with the TotalSeq-C antibody mix: TotalSeq - C0048 anti-human CD45 Antibody (Biolegend; Cat# 368545), TotalSeq - C0103 anti-mouse/human CD45R/B220 (Biolegend; Cat# 103273), TotalSeq - C0087 anti-human CD45RO (Biolegend; Cat# 304259), and TotalSeq - C0063 anti-human CD45RA (Biolegend; Cat# 304163). Before adding the surface antibody mix, equal volumes of each antibody were combined and the mix was spun at 14,000rpm for 5min to remove aggregates. After staining the cells were washed twice with staining buffer, and a final wash was completed in DMEM with 10% FCS before counting. Single-cell RNA libraries were generated using the 10x Genomics Chromium Single Cell V(D)J Reagent Kit using 5’ v1 chemistry with Feature Barcode technology for Cell Surface Protein (10x Genomics; Cat# 1000080). After each step, cDNA generation, gene expression libraries, and cell surface protein libraries samples quality was assessed using the Qubit dsDNA high sensitivity kit (Invitrogen; Cat# Q32854) and the high sensitivity BioA DNA kit (Agilent; Cat# 5067-4626). Samples that passed quality control were sequenced on a NextSeq 500 sequencer (Illumina), using pair-end reads, with 26 reads for read 1 and 55 reads for read 2.

### Multiplexed array assembly of cDNA libraries

cDNA libraries were amplified using the following reaction conditions: 34μL of H2O, 50μL of Kapa HiFi Uracil+ ReadyMix (2X) (Roche #7959079001), 5μL of primer AAO272 (10μM, IDT), 5μL of primer AAO273 (10μM, IDT), and 6μL 10x 5’ cDNA library (∼3ng/μL) and the following cycling conditions: 98 °C for 3 min, followed by 5 cycles of 98 °C for 20 s, 65 °C for 30 s and 72 °C for 8 min, followed by a final 72 °C extension for 10 min. Amplified libraries were purified using 0.7x SPRIselect (Beckman Coulter B23318) cleanup and quantified using Qubit (Thermo #Q32851). Libraries were further purified using 10μL (100μg) Dynabeads™ kilobaseBINDER™ (Thermo #60101) with final bead reconstitution in 40μL TE (Thermo #AM9849) after binding/washing. After streptavidin purification, 2ul of USER^®^ Enzyme (M5505S) was added and incubated at 37 °C for 2 hours to uncouple the bound cDNAs from the beads. Following USER digestion, the reaction was placed on a magnet for 5 minutes, separating the beads and supernatant containing the cDNAs. The cDNA fraction was moved to a fresh tube and purified using 0.7x SPRIselect (Beckman Coulter B23318) cleanup. After cDNA purification, the following PCR master mix was assembled: 580μL of H2O, 750μL of Kapa HiFi Uracil+ ReadyMix (2X) (Roche #7959079001), and 20μL 10x 5’ cDNA library (∼6ng/μL). 90μL of the mastermix was distributed in 15 PCR tubes, each containing 10μL of 5μM MAS-seq primer pair mix. The 15 reactions were then thermocycled with the following cycling conditions: 98 °C for 3 min, followed by 8 cycles of 98 °C for 20 s, 65 °C for 30 s and 72 °C for 8 min, followed by a final 72 °C extension for 10 min (optimal cycling number was identified using scaled down qPCR reaction). Reactions were then pooled in a 5 ml tube and purified using a 0.7x SPRIselect (Beckman Coulter B23318) cleanup and eluted in 450μL of TE. In a subsequent reaction, 15μL of USER^®^ Enzyme (M5505S) was added to 435μL of the pooled product and set to incubate at 37 °C for 2 hours. Following USER digestion, 15 μL HiFi *Taq* DNA Ligase (M0647S) and 51 μL of HiFi *Taq* DNA Ligase buffer was added to the reaction and incubated in a thermocycler at 42 °C for 2 hours. Following ligation, the reaction was purified using a 0.7x AMPure PB Bead (Pacific Biosciences #100-265-90) cleanup and eluted in 180μL of H2O. Multiplexed array libraries were quantified using Qubit (Thermo #Q32851) and Genomic DNA ScreenTape (Agilent #5067-5365).

### SIRV-Set 4 cDNA generation

SIRV-Set 4 (Lexogen #141.01) was thawed and aliquoted 1μL into each of 9 PCR tubes on ice. Following primary aliquoting, 2ul of Tris-EDTA pH7.0 was added to each tube and mixed. SIRV stocks were then frozen at -80 °C. For first strand synthesis, the following primary master-mix was set up: 15.5μL of H2O, 3.2μL of Polyethylene glycol 8,000 50% (w/v) (VWR #25322-68-3), 0.24μL Triton X-100 10% solution (Fisher #9002-93-1), 0.32μL SUPERase•In™ RNase Inhibitor (Thermo #AM2696), 1.6μL of dNTP mix 10mM (NEB #N0447S), 0.16μL OligodT primer 100μM (IDT) (SS3_OligodTVN for Smart-seq3 and MAS_OligodTVN for Iso-seq and MAS-seq), 3μL of SIRV-Set 4 aliquot. Additionally, the following RT master-mix was assembled: 1.2μL of H2O, 0.8μL of Tris-HCl pH 8.5 (1M), 0.96μL of NaCl (1M), 0.8μL of MgCl2 (100mM), 0.32μL of GTP (100mM), 2.56μL of DTT (100mM), 0.4μL of SUPERase•In™ RNase Inhibitor (Thermo #AM2696), 0.64μL of TSO 100uM (IDT) (SS3_OligodTVN for Smart-seq3 and MAS_OligodTVN for Iso-seq and MAS-seq), 0.32μL of Maxima H-minus RT enzyme 200U/uL (Thermo #EP0751). Both primary and RT master-mixes were added to the thermocycler with the following conditions: 42 °C for 90 min, followed by 10 cycles of 50 °C for 2 min, 42 °C for 2 min, followed by a final 85 °C 5 min.

### Smart-seq3 of SIRV-Set 4

To amplify the cDNA, the cDNA generation reaction was added to straight into the following PCR mix: 26.5μL of H2O, 16μL of Kapa HiFi HotStart buffer (5X), 2.4μL of dNTP mix 10mM (NEB #N0447S), 0.4μL of MgCl2 (100mM), 0.4μL of fwd_primer 100μM (IDT), 0.8μL of rev_primer 10μM (IDT), 1.6μL of Kapa Hifi DNA polymerase (KK2103). The reaction was amplified using the following conditions: 98 °C for 3 min, followed by 13 cycles of 98 °C for 20 s, 65 °C for 30 s and 72 °C for 8 min, followed by a final 72 °C extension for 10 min. Amplified cDNA libraries were purified using 0.7x SPRIselect (Beckman Coulter B23318) cleanup and quantified using Qubit (Thermo #Q32851). Libraries were normalized to 0.1ng/μL and tagmented using the following reaction conditions: 7.56μL of H2O, 9μL of Tagmentation buffer 4x (Tris-HCl pH 7.5 (40mM), MgCl2 (20mM), DMF (20%)), 1.44μL Amplicon Tagmentation Mix (XYZ), 4μL of normalized cDNA libraries. Tagmentation reaction was mixed, spun down, then added to a thermocycler at 55 °C for 10 min. After tagmentation, 2μL of 2% SDS was immediately added and incubated for 5 min to halt the reaction. To the tagmented cDNA reactions, 6μL of nextera primer pair mixes (0.5μM) were added. Following addition of primers, the following PCR was assembled: 25.38 μL of H2O, 25.2μL of Phusion Buffer 5x (Thermo Scientific #F530L), 2.7μL of dNTP mix 10mM (NEB #N0447S), 0.72μL of Phusion High-Fidelity DNA Polymerase 2 U/μL and added to the thermocycler with the following conditions: 72 °C for 3 min, 98 °C for 3 min, followed by 12ncycles of 98 °C for 10 s, 55 °C for 30 s and 72 °C for 30 s, followed by a final 72 °C extension for 5 min. Amplified final libraries were purified using 0.7x SPRIselect (Beckman Coulter B23318) cleanup and quantified using Qubit (Thermo #Q32851) and Agilent High Sensitivity DNA kit for BioAnalyzer (Agilent #5067-4626). Libraries were sequenced on an Illumina NovaSeq 6000, using paired-end 150 read lengths.

### Smart-seq3 processing workflow

#### Aligning and stitching UMI-containing reads

We process Smart-seq3 SIRV Illumina paired-end reads closely following the procedure outlined in Ref. ^6^. We processed raw non-demultiplexed FASTQ files using zUMIs v2.9.4g and STAR v2.5.4b in order to generate expression profiles for both the 5′ UMI-containing and internal reads. To extract and identify the UMI-containing reads in zUMIs, we specified find_pattern: ATTGCGCAATG for the 5’ read together with base_definition: cDNA (23–150), UMI (12–19) in the configuration YAML file and collapsed UMIs within a Hamming distance of 1. Next, we stitched UMI-containing reads together using stitcher.py^16^ starting from the <prefix>.filtered.Aligned.GeneTagged.UBcorrected.sorted.bam output from zUMIs. We inferred the transcript compatibility set for each 5′ UMI-containing read from the CT tag in the produced BAM file. Finally, we generated the transcript identification confusion matrix by iterating over all stitched 5’ reads, assuming a flat prior for both source and target transcripts, and accordingly dividing the assignment probability weight equally to all compatible source and target transcripts.

### Quantification of SIRV isoforms

We quantified the abundance of Smart-seq3 SIRV isoforms from both 5’ UMI-containing and internal reads. To this end, we ran salmon v1.5.1 in quantification mode with additional arguments “--minAssignedFrags 1 -l IU” on the previously obtained <prefix>.filtered.tagged.Aligned.toTranscriptome.out.bam transcriptome alignments from zUMIs. We read the TPM normalized abundances from the salmon_quant/quant.sf output table.

### MAS-seq processing workflow

#### Error correction

Error correction was performed on-board the PacBio Sequel IIe with the vendor’s ccs software v5.0.0^7^ and settings “--all --subread-fallback --num-threads 232 --streamed <movie_name>.consensusreadset.xml --bam <movie_name>.reads.bam”. With these settings, all reads from the instrument (including those failing ccs correction) are presented in a single BAM^17^ file for downstream analysis. Each read is affixed with an auxiliary BAM tag “rq” indicating overall read quality ranging from rq=-1 for uncorrected reads, -1 < rq < 0.99 for corrected reads with predicted accuracy < Q20, and rq ≥ 0.99 for corrected reads with predicted accuracy ≥ Q20^18^.

#### Annotation/filtration/segmentation/demultiplexing

We developed a composite hidden Markov model (“*Longbow*”) to enable the per-read labeling of all subsequences of interest (annotation), allowing for insertions, deletions, and mismatches in both low and high error rate data. Given a predefined array structure, we combined several instances of two probabilistic models for pairwise sequence alignment: the Needleman-Wunsch and random alignment models^19^. Needleman-Wunsch model sections support annotation of sequences known *a priori* (i.e. MAS-seq adapters; 10x Genomics single-cell 5’ and 3’ adapters; poly(A) tails). Random alignment model sections support annotation of unknown interstitial sequences (i.e. cDNA sequences) and barcode identifiers. In this formulation, a MAS-ISO-seq read is considered to be a mosaic of imperfect (but complete) copies of the various known adapter sequences among which the unknown cDNA sequences of interest are present.

The state transition diagram and default values for transmission and emission probabilities (used for all MAS-ISO-seq processing performed in this work) are provided in Supplementary Fig. 14. These defaults can optionally be refined using *Longbow’s* train command, which will estimate the parameters of the model using Baum-Welch learning.

Data processing proceeds as follows: annotations as described above are generated for both the forward and reverse-complement orientations, retaining the result from the model with higher log-likelihood. Given the design expectation that MAS-ISO-seq adapters should be found in sequence along the length of the read, we verify that each read conforms to this expectation and filter out any read with mis-ordered MAS-ISO-seq adapters. We then segment each read between MAS-ISO-seq adapters and 10x Genomics single-cell 5’ adapter.

For multiplexed libraries (e.g. libraries with different array configurations and run on the same flow cell), the demultiplexing workflow proceeds similarly to the procedure described above with one notable change: annotations are generated for both the forward and reverse-complement read orientations and over each user-specified array design. The annotations from the read orientation and array design that maximize the overall log-likelihood are propagated to subsequent steps.

### Cell Barcode (CBC) and UMI Annotation

We annotated segmented reads with CBC and UMIs by leveraging the structure of the read library. Each read begins with a 12 base MAS-seq adapter, 22 bp forward adapter, a 16 bp cell barcode, then a 10 bp UMI. We aligned the forward adapter to the 200 bases on either end of each read using an accelerated Smith-Waterman algorithm, SSW (v1.2.4)^20^, to determine a known position in the read, then counted bases from the end of that alignment to annotate each read with the raw CBC and UMI. We also annotated each read with the base qualities for the CBC (found using the same offsets). In the case of SIRV data, no CBC was present in the library and therefore it was not annotated.

### CBC Error Correction

To ensure single-cell data analysis could be performed accurately, we performed CBC error correction by integrating evidence from multiple data sources according to the following procedure. We first annotated each long read that passed *Longbow* filtering with a raw CBC. We further sequenced the T cell samples on Illumina instruments and processed the data with Cell Ranger v3.1.0^21^ to annotate each short read with a cell barcode and the corresponding base qualities. For each read, including long CCS-corrected, long CCS-uncorrected, and short (Illumina), we computed a barcode pseudocount (BPC) from the barcode sequence base qualities as follows:

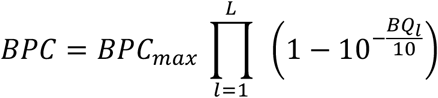

where *BQ*_*l*_ is the Phred-scaled base quality of the barcode letter at position *l*, and *L* = 16 is the barcode length. The prefactor *BPC*_*max*_ = 100 is arbitrary and is for reducing quantization noise in rounding the aggregated pseudocounts to whole integers (see below). Intuitively, BPC is proportional to the probability for the observed barcode sequence to coincide with the actual barcode sequence. BPC is small for low-quality barcode sequences, e.g. CCS-uncorrected reads, and reaches ∼ *BPC*_*max*_ for high-confidence observations, e.g. Illumina reads or CCS-corrected reads. We combined the pseudocounts from all available reads for each of the observed barcode sequences, rounded the resulting counts to whole integers, and used Starcode v1.4 2020-11-02^20,22^ to cluster the cell barcodes using aggregated pseudocounts in place of a raw counts (starcode --print-clusters --quiet -i). Finally, we used the starcode output to create a mapping from raw cell barcodes to corrected cell barcodes (i.e. cluster centroids), and annotated each read in our long-read datasets with the corresponding corrected cell barcode. Notably, error correction can be performed without a whitelist or additional Illumina sequencing (in which case, the CBC from high-confidence CCS-corrected reads will drive the CBC error correction of noisy CCS-uncorrected reads).

### Quantification of SIRV isoforms

To quantify SIRV isoforms, we took reads (both CCS-corrected and CCS-uncorrected) that had been filtered, annotated, and segmented by *Longbow* and annotated their UMIs. We then removed the adapter sequences and poly-A tails from these reads. The resulting reads were aligned to the SIRV-Set 4 transcriptome using minimap2 v2.17-r941^23^ with the HiFi read preset (minimap2 -ayYL --MD --eqx -x asm20). We then took the primary alignments, removed any in which we could not detect a UMI, and annotated each one with the contig on which they were aligned (in this case, a transcript). These UMI-containing primary alignments were then processed using UMI-tools v1.1.1^24^ to group the detected UMIs and mark the molecules with the same UMIs (umi_tools group --buffer-whole-contig --no-sort-output --per-gene --gene-tag XG --extract-umi-method tag --umi-tag ZU --group-out=out.tsv --output-bam). Finally, we created a count matrix from the resulting BAM file by tallying for each transcript and CBC how many unique UMI occurrences there were.

### Quantification of 10x Genomics 5’ CD8+ T cell isoform expression

To quantify T cell isoforms, we took reads, both CCS-corrected and CCS-uncorrected, that had been filtered, annotated, and segmented by *Longbow* and annotated the CBCs and UMIs contained in each. We corrected the CBC in each read and removed the adapter sequences and poly-A tails. These resulting extracted reads were aligned to a version of the GRCh38 human reference genome with alternate contigs removed using minimap2 v2.17-r941 with the splicing preset (minimap2 -ayYL --MD --eqx -x splice:hq). We then processed the resulting reads with StringTie2 v2.1.6^25^ using Gencode v37 as baseline transcript annotations to create new transcriptome annotations specific to each of our samples (stringtie -Lv -G gencode.v37.primary_assembly.annotation.gtf -o annotations.gtf -A gene_abund.out). Based on the resulting transcript annotations we created a new transcript reference FASTA file using gffread v0.12.6 (gffread -w transcriptome.fa -g hg38_no_alt.fasta annotations.gtf). We then aligned the extracted reads to the novel transcriptome reference using minimap2 v2.17-r941 (minimap2 -ayYL --MD --eqx -x asm20). We then took the resulting primary alignments, removed any in which we could not detect a UMI, and annotated each one with the contig on which they were aligned (in this case, a transcript). These UMI-containing primary alignments were then processed using UMI-tools v1.1.1 to group the detected UMIs and mark the molecules with the same UMIs (umi_tools group --buffer-whole-contig --no-sort-output --per-gene --gene-tag XG --extract-umi-method tag --umi-tag ZU --group-out=out.tsv --output-bam). We then created a count matrix from the resulting bam file by tallying for each transcript and CBC combination how many unique UMI occurrences there were.

### Single cell analysis

#### Short-read 10x Genomics 5’ gene expression and antibody capture preprocessing

We quantified the produced 5’ RNA capture and TotalSeq-C antibody capture libraries using Cell Ranger v3.1.0 count workflow. We imported the count data into AnnData format using scanpy v1.7.2 read_10x_h5 command. Our preliminary investigations indicated that the Cell Ranger automatic cell identification algorithm had used an excessively conservative cutoff, leading to the loss of 30%-50% viable non-empty droplets (primarily of stem-like memory T cells origin, a cell type that exhibits relatively lower transcriptional complexity). As a countermeasure, we loaded the raw count data from Cell Ranger count output raw_feature_bc_matrix.h5 and kept every droplet expressing > 500 unique genes and > 80% non-mitochondrial genes. We performed a preliminary round of clustering and differential gene expression analysis using scanpy standard workflow^26^. We identified and removed non-immune cell clusters of likely primary tumor origin. We additionally identified and removed doublets using scrublet v0.2.3. The estimated doublet rate was 14%, which is the expected figure for loading ∼10,000 cells. Finally, we log-transformed the antibody capture counts and treated them as cell-level annotations for the rest of the analysis.

### Long-read single-cell MAS-ISO-seq isoform expression preprocessing

As a first step, we converted the isoform-level UMI count matrix produced by the MAS-ISO-seq workflow to an AnnData object. During this conversion, additional metadata was added to the transcript counts. Many of the novel transcripts and genes discovered by StringTie2 could be unambiguously assigned back to a GENCODE annotation. In particular, novel genes were assigned to known genes in GENCODE v37 if the novel genes had transcripts overlapping exactly one unique gene in GENCODE v37. Novel genes with transcripts overlapping multiple GENCODE genes were marked as ambiguous and removed from the analysis. In addition, an interval list containing T cell receptor genes^27^ was cross-referenced and transcripts found overlapping these intervals were accordingly marked. To harmonize the long- and short-read AnnData objects for joint analyses, we only kept the mutual cell barcodes between the two datasets. Remarkably, we could identify > 98.8% of T cell barcodes identified from the short-read dataset in the MAS-ISO-seq long-read dataset, indicating the high fidelity of our CBC error correction algorithm.

### Normalization, clustering, and embedding

We imported the harmonized short- and long-read AnnData objects to seurat v4.0.3 using SeuratData v0.2.1 and SeuratDisk v0.0.0.9015 helper packages^28^. We performed a negative binomial (NB) variance-stabilizing transformation (VST) on each count dataset separately using sctransform v0.3.2. We treated isoform counts similarly to gene counts, which is justified since isoform counts exhibit the same class of technical noise and statistical dropout as gene counts. Given the much larger number of isoforms, we found it necessary to increase the number of isoforms used for training the NB model from the default value of 2,000 to 10,000. We did not notice any significant change in the downstream results by increasing this figure any further. The Pearson residuals for all cells and genes were exported to AnnData. We performed clustering and embedding separately for short- and long-read datasets using the same workflow as follows. We selected the top 5,000 genes (or isoforms) sorted in the descending order of total Pearson residual as highly variable features (HVF). The HVFs were z-scored independently to equalize the role of each gene (or isoform). We reduced the feature set down to 30 using PCA and calculated the k=100 nearest neighbor graph for each cell in the PCA space based on the Euclidean distance. The resultant neighbor graph was used for obtaining a 2D embedding using UMAP, and clustering using the Leiden algorithm with resolution parameter set to 1.1. We performed differential gene expression (DE) analysis on the short-read dataset based on *t*-test, which is an appropriate statistical test for VST counts, as implemented in scanpy rank_genes_groups method. The DE genes were used for annotating the clusters shown in Fig. 2 using known T cell subtype markers.

### Diffusion pseudotime analysis

We performed diffusion pseudotime (DPT) analysis closely following the scanpy hematopoiesis trajectory analysis workflow^29^ with one notable modification. We noticed that using scaled highly-variable Pearson residuals in place of log-transformed counts resulted in significantly cleaner force-directed graphs. The latter is expected given that Pearson residuals are more Gaussian-like compared to log-transformed counts, and thus, better suited to the assumptions of the DPT model. Accordingly, we substituted the standard preprocessing and normalization step with the sctransform workflow.

### CD45 isoform annotation refinement

The GENCODE v37 contains a rather extensive set of isoform annotations for CD45, including the RO, RABC, RB, RBC, and RAB. The annotations, however, are frequently incomplete and miss a large portion of the coding sequences. For instance, out of the available annotations for *PTPRC* (CD45), only two (ENST00000348564: CD45RO and ENST00000442510: CD45RABC) extend all the way to the 3’ UTR (Supplementary Fig. 15). Given the short-reads origin of currently available transcriptome annotations, we expect this caveat to prevail among most other genes, as also indicated by other authors^30,31^. In particular, we noticed that the more complete and longer annotations, such as RO and RABC, tend to be preferred primary target by the aligner (minimap2) for the vast majority of reads, overriding the differential usage of short but biologically important exons such as the A, B, and C. We found it advantageous to refine GENCODE annotations using StringTie2, which resulted in the extension of several incomplete GENCODE annotations and significantly improved the specificity of isoform assignments to different CD8+ T cell subtypes (Supplementary Fig. 15). Ultimately, though, we found the refined annotations produced by StringTie2 to be still incomplete and direct transcriptome alignment to be lacking in specificity. Inspired by the highly distinct structure of the CD45 sashimi plots for different T cell subtypes (Supplementary Fig. 15), we reasoned higher specificity could be achieved by directly classifying genomics alignments. To this end, we mapped the reads to the genome (GRCh38, alternate contigs removed) using minimap2 v2.17-r941 with the splicing preset (minimap2 -ayYL --MD --eqx -x splice:hq), and performed CD45 isoform assignment using a decision tree based on the presence/absence of landmark exons, e.g. A, B, and C in CD45 (Supplementary Fig. 16; note the increased specificity of isoform assignment to different T cell subtypes). The results shown in Fig. 2b, c are based on the decision tree methodology. Leveraging the significantly increased read depth afforded by MAS-ISO-seq, we find improving algorithms for *de novo* isoform identification and clustering, and benchmarking the available isoform quantification pipelines (e.g. FLAIR^31^, TALON^32^, etc.), and producing more complete transcriptome annotation references to be crucial areas of future method and resource development.

### Identification of differentially spliced genes

We consider two types of differential splicing (DS) statistical tests for every expressed gene. *(global DS test)* First, we wish to determine whether the isoforms of a given gene are differentially expressed in different cell clusters. To this end, we produce a contingency table with isoforms and cell clusters as rows and columns, and with the aggregated isoform expression counts as entries. A non-trivial global DS pattern is equivalent to having a statistical dependence between the columns and rows of this contingency table. The latter, however, can be canonically assessed using Fisher’s exact test generalized to arbitrary *m* × *n* contingency tables with *m, n* ≥ 2. Notably, we found the requirements for fast Chi-squared asymptotic approximation to be out of reach for the majority of cases. Therefore, we use the fisher.test as implemented in R v4.1.1 to perform the test using 1*e*6 permutations. *(cluster-resolved DS test)* We additionally perform a cluster-resolved DS test for every gene, whereby we wish to know whether a gene exhibits differential isoform usage in each of the clusters vs. the rest. Like before, we form a contingency table with two columns signifying the cluster of interest and the rest, isoforms in rows, and aggregated isoform expression counts as entries. We similarly obtain a p-value by performing a permutation-based Fisher’s test for every gene and every cluster. Finally, for both tests, we treat the obtained p-values as a collection of independent hypotheses and adjust the p-values for false discovery rate (FDR) at level α = 0.05 using the Benjamini-Hochberg step-up procedure.

### Table of MAS-seq barcode adapters

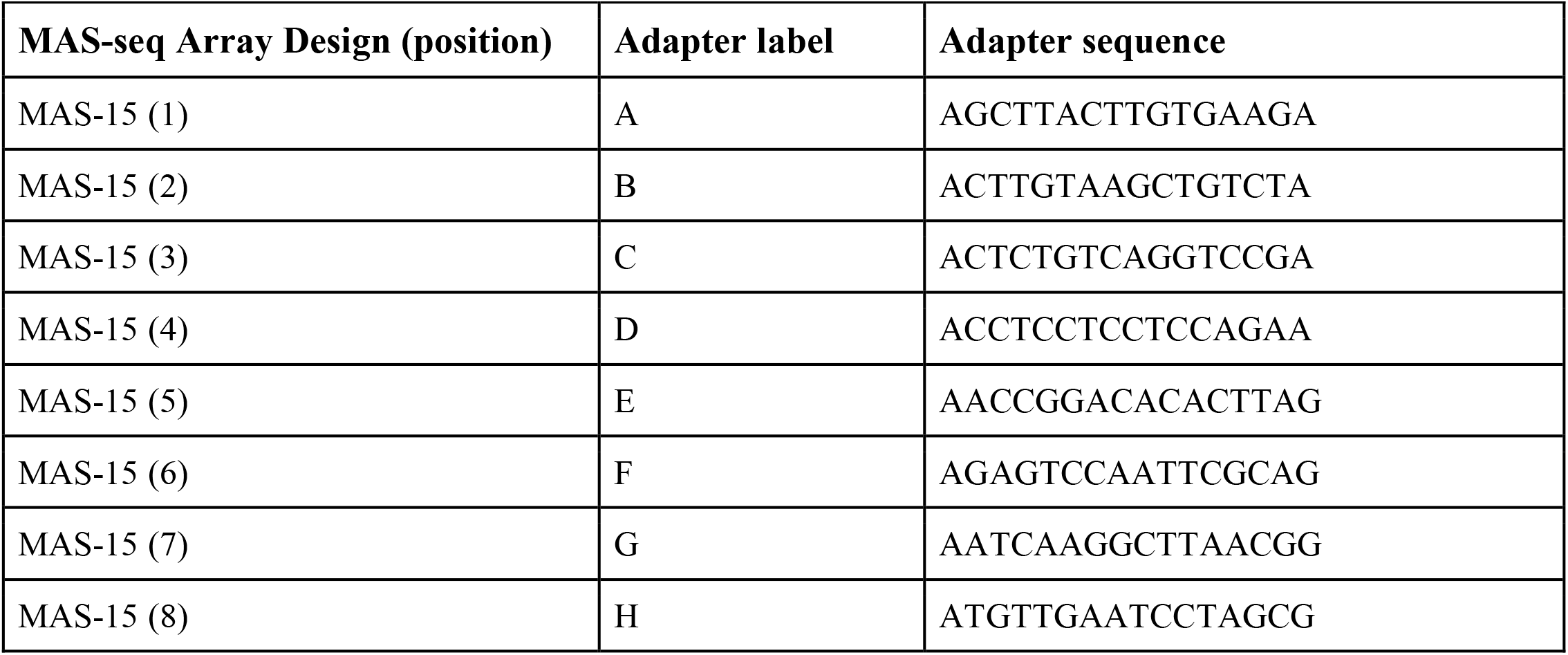
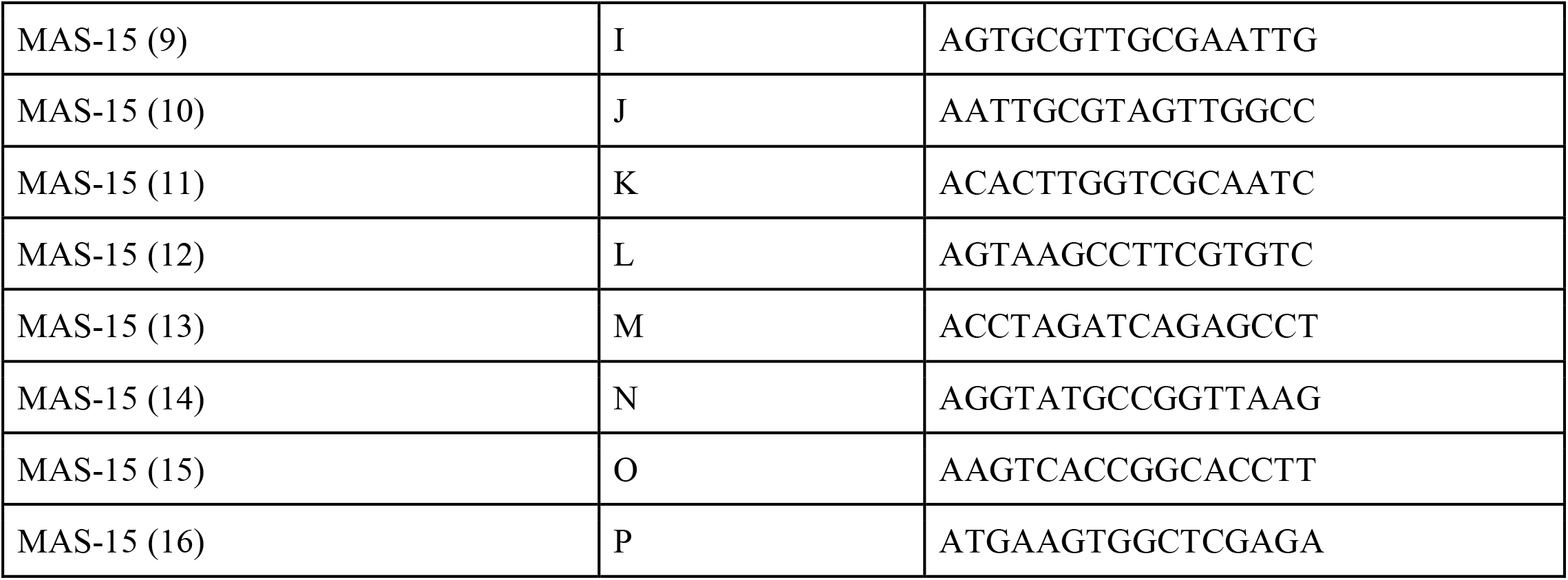

### Table of MAS-seq primers

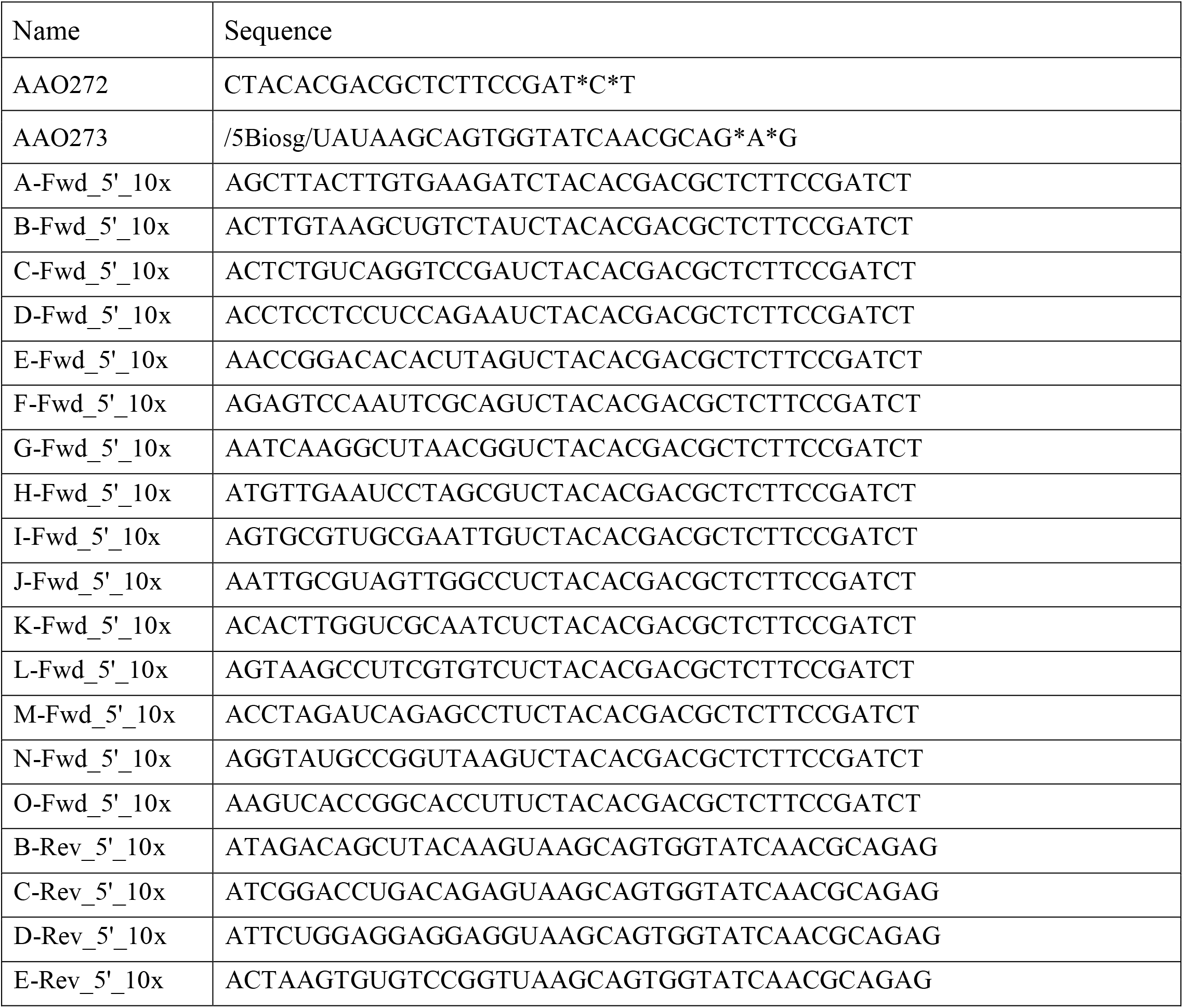
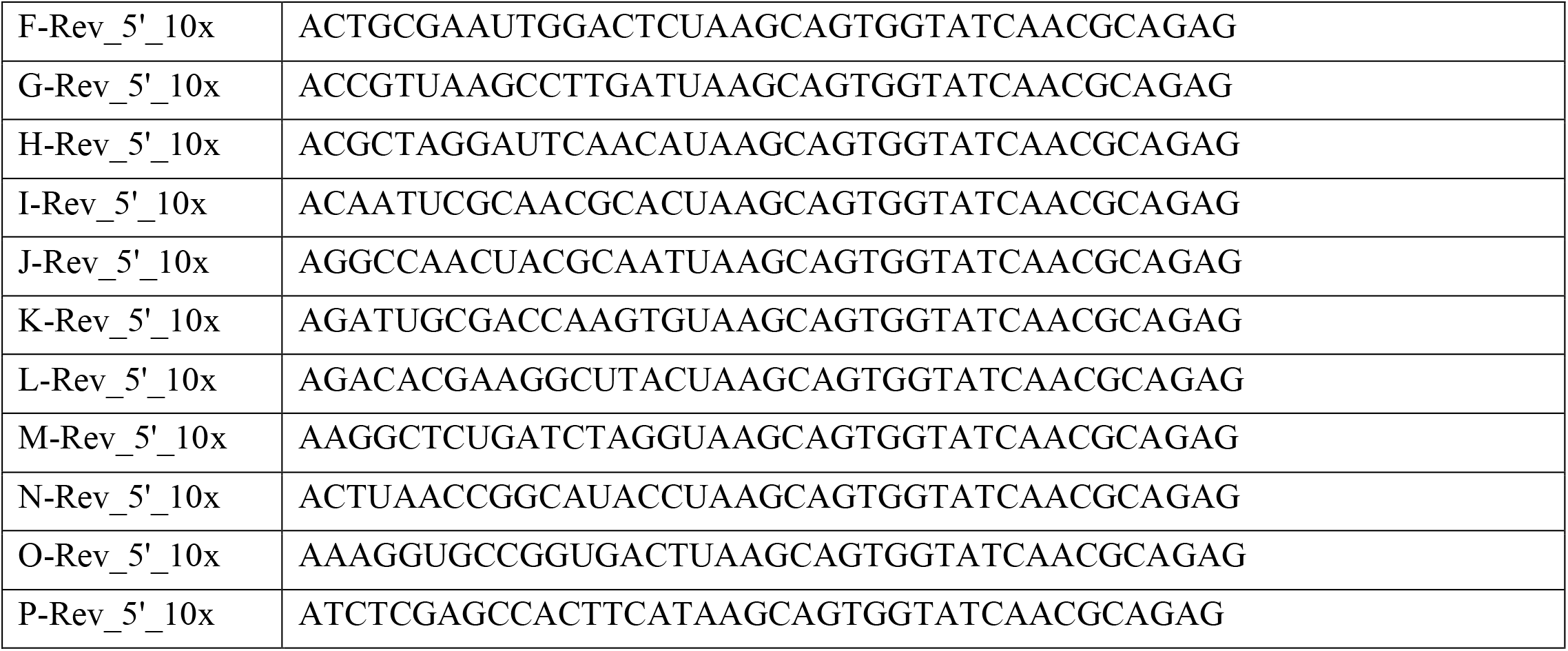

## Data Availability

Links to the datasets used in this study can be found at https://github.com/broadinstitute/mas-seq-paper-data.

## Code Availability

An online repository of code used in this study can be found at https://github.com/broadinstitute/longbow.

## Acknowledgments

We thank W. Kretzschmar for helpful discussions. This work was supported by Broad Institute SPARC awards #800353 (A.M.A., K.V.G., N.H., and P.B.) and #800307 (K.V.G.); National Institutes of Health grants U19 AI082630 (N.H.), Adelson Medical Research Foundation (N.H.), RM1HG006193 (N.H., P.C.B.), with additional support from the Center for Cell Circuits at the Broad Institute (HG006193). M.A.S. is a Cancer Research Institute Irvington Fellow supported by the Cancer Research Institute (CRI Award 4071).

## Author information

These authors contributed equally: Aziz Al’Khafaji, Jonathan T. Smith, Kiran V Garimella, Mehrtash Babadi.

**Maura Costello**

Present affiliation: seqWell, Beverly, MA, USA

Affiliations

**Cell Circuits, Broad Institute of MIT and Harvard, Cambridge, MA, USA**

Aziz Al’Khafaji, Emily M. Blaum, Siranush Sarkizova, Nir Hacohen, Paul Blainey

**Data Sciences Platform, Broad Institute of MIT and Harvard, Cambridge, MA, USA**

Jonathan T. Smith, Kiran V Garimella, Mehrtash Babadi, Michael Gatzen, Anthony A Philippakis, Eric Banks

**Broad Institute of MIT and Harvard, Cambridge, MA, USA**

Victoria Popic

**Genomics Platform, Broad Institute of MIT and Harvard, Cambridge MA, USA**

Maura Costello, Allyson Day, Tera Bowers, Stacey Gabriel

**Department of Medicine, Center for Cancer Research, Massachusetts General Hospital, Boston, MA, USA**

Moshe Sade-Feldman, Nir Hacohen

**Department of Biological Engineering, Massachusetts Institute of Technology, Cambridge, MA, USA**

Paul Blainey

**Koch Institute for Integrative Cancer Research at Massachusetts Institute of Technology, Cambridge, MA, USA**

Paul Blainey

**Harvard Medical School, Boston, MA, USA**

Moshe Sade-Feldman, Marc A. Schwartz, Nir Hacohen

**Department of Surgery, Massachusetts General Hospital**

Genevieve M. Boland

## Contributions

A.M.A. conceived of and developed the DNA concatenation workflow and designed and performed the experiments. K.V.G. developed the statistical annotation software. J.S. developed the data processing pipeline with contributions from M.G., and performed bioinformatic analyses. M.B. performed Smart-seq3 analysis, scRNA-seq data analysis and statistical modeling, and devised the barcode error correction algorithm. V.P. evaluated alternative isoform assignment algorithms. S.S. aided in computational analysis and discussion. G.M.B., E.M.B., and M.S.F. consented patients, collected samples, processed, and generated the 10x Genomics scRNA-seq data. M.A.S assisted with T cell data analysis. M.C., A.D., T.B., and S.G. aided in the data generation and helped troubleshoot early iterations of the protocol. A.A.P. and E.B. provided access to cloud compute and other resources to facilitate data processing and analysis. A.M.A., K.V.G., J.S., M.B., P.B., and N.H. co-wrote the manuscript.

## Ethics declarations

### Competing interests

A.M.A., K.V.G., J.S., M.B., P.B., and N.H. have filed a patent on the MAS-seq method.

A.A.P. is a Venture Partner and Employee of GV. He has received funding from Verily, Microsoft, Illumina, Bayer, Pfizer, Biogen, Abbvie, Intel, and IBM.

M.S.F. receives funding from Bristol-Myers Squibb.

G.M.B. has served on SAB and on the steering committee for Nektar Therapeutics. She has SRAs with Olink proteomics and Palleon Pharmaceuticals. She served on SAB and as a speaker for Novartis

N.H. holds equity in BioNTech and is a founder and equity holder of Danger Bio.

P.C.B. is a consultant to and/or holds equity in companies that develop or apply genomic or genome editing technologies: 10X Genomics, General Automation Lab Technologies, Celsius Therapeutics, Next Gen Diagnostics LLC, Cache DNA, and Concerto Biosciences. P.C.B. receives funding from industry for unrelated work.

## References

1. Hardwick, S. A., Joglekar, A., Flicek, P., Frankish, A. & Tilgner, H. U. Getting the Entire Message: Progress in Isoform Sequencing. Front. Genet. 10, 709 (2019).

2. Baralle, F. E. & Giudice, J. Alternative splicing as a regulator of development and tissue identity. Nat. Rev. Mol. Cell Biol. 18, 437–451 (2017).

3. Scotti, M. M. & Swanson, M. S. RNA mis-splicing in disease. Nat. Rev. Genet. 17, 19–32 (2015).

4. Dvinge, H., Kim, E., Abdel-Wahab, O. & Bradley, R. K. RNA splicing factors as oncoproteins and tumour suppressors. Nat. Rev. Cancer 16, (2016).

5. Kanitz, A. et al. Comparative assessment of methods for the computational inference of transcript isoform abundance from RNA-seq data. Genome Biol. 16, 1–26 (2015).

6. Hagemann-Jensen, M. et al. Single-cell RNA counting at allele and isoform resolution using Smart-seq3. Nat. Biotechnol. 38, 708–714 (2020).

7. Wenger, A. M. et al. Accurate circular consensus long-read sequencing improves variant detection and assembly of a human genome. Nat. Biotechnol. 37, 1155–1162 (2019).

8. Volden, R. et al. Improving nanopore read accuracy with the R2C2 method enables the sequencing of highly multiplexed full-length single-cell cDNA. Proc. Natl. Acad. Sci. U. S. A. 115, 9726–9731 (2018).

9. Tilo Buschmann <tilo. buschmann. >. DNABarcodes. (Bioconductor, 2017). doi:10.18129/B9.BIOC.DNABARCODES.

10. Paul, L. et al. SIRVs: Spike-In RNA Variants as External Isoform Controls in RNA-Sequencing. bioRxiv 080747 (2016) doi:10.1101/080747.

11. Oberdoerffer, S. et al. Regulation of CD45 Alternative Splicing by Heterogeneous Ribonucleoprotein, hnRNPLL. Science 321, 686–691 (2008).

12. Bio-Rad. CD45 characterization & Isoforms - Mini-review. https://www.bio-rad-antibodies.com/cd45-characterization-isoforms-structure-function-antibodies-minireview.html.

13. Zheng, Y.-F. et al. HIT-scISOseq: High-throughput and High-accuracy Single-cell Full-length Isoform Sequencing for Corneal Epithelium. bioRxiv 2020.07.27.222349 (2020) doi:10.1101/2020.07.27.222349.

14. Schlecht, U., Mok, J., Dallett, C. & Berka, J. ConcatSeq: A method for increasing throughput of single molecule sequencing by concatenating short DNA fragments. Sci. Rep. 7, 1–10 (2017).

15. Kanwar, N., Blanco, C., Chen, I. A. & Seelig, B. PacBio sequencing output increased through uniform and directional fivefold concatenation. Sci. Rep. 11, 1–13 (2021).

16. Larsson, A. J. M. & Sandberg, R. stitcher.py. (2020). doi:10.5281/zenodo.3765223.

17. Li, H. et al. The Sequence Alignment/Map format and SAMtools. Bioinformatics 25, 2078 (2009).

18. Pacific Biosciences, Inc. What is in the reads.bam? CCS Docs https://ccs.how/faq/reads-bam.html.

19. Durbin, R., Eddy, S. R., Krogh, A. & Mitchison, G. Biological Sequence Analysis. (1998) doi:10.1017/cbo9780511790492.

20. Zhao, M., Lee, W.-P., Garrison, E. P. & Marth, G. T. SSW Library: An SIMD Smith-Waterman C/C++ Library for Use in Genomic Applications. PLoS One 8, e82138 (2013).

21. Zheng, G. X. Y. et al. Massively parallel digital transcriptional profiling of single cells. Nat. Commun. 8, 1–12 (2017).

22. Zorita, E., Cuscó, P. & Filion, G. J. Starcode: sequence clustering based on all-pairs search. Bioinformatics 31, 1913–1919 (2015).

23. Li, H. Minimap2: pairwise alignment for nucleotide sequences. Bioinformatics 34, 3094–3100 (2018).

24. Smith, T. S., Heger, A. & Sudbery, I. UMI-tools: Modelling sequencing errors in Unique Molecular Identifiers to improve quantification accuracy. Genome Res. gr.209601.116 (2017).

25. Kovaka, S. et al. Transcriptome assembly from long-read RNA-seq alignments with StringTie2. Genome Biol. 20, 1–13 (2019).

26. Alex Wolf, Fidel Ramirez, Sergei Rybakov. Preprocessing and clustering 3k PBMCs. Scanpy documentation https://scanpy-tutorials.readthedocs.io/en/latest/pbmc3k.html.

27. HGNC. Gene group: T cell receptors (TR). HUGO Gene Nomenclature Committee https://www.genenames.org/data/genegroup/#!/group/370.

28. Hao, Y. et al. Integrated analysis of multimodal single-cell data. Cell 184, 3573–3587.e29 (2021).

29. Alex Wolf, Fidel Ramirez, Sergei Rybakov. Trajectory inference for hematopoiesis in mouse. Scanpy documentation https://scanpy-tutorials.readthedocs.io/en/latest/paga-paul15.html.

30. Glinos, D. A. et al. Transcriptome variation in human tissues revealed by long-read sequencing. bioRxiv 2021.01.22.427687 (2021) doi:10.1101/2021.01.22.427687.

31. Tang, A. D. et al. Full-length transcript characterization of SF3B1 mutation in chronic lymphocytic leukemia reveals downregulation of retained introns. Nat. Commun. 11, 1–12 (2020).

32. Seki, M., Oka, M., Xu, L., Suzuki, A. & Suzuki, Y. Transcript Identification Through Long-Read Sequencing. Methods Mol. Biol. 2284, 531–541 (2021).

