## Supplementary Information for "High-throughput RNA isoform sequencing using programmable cDNA concatenation"

* - These authors contributed equally

† - Corresponding authors

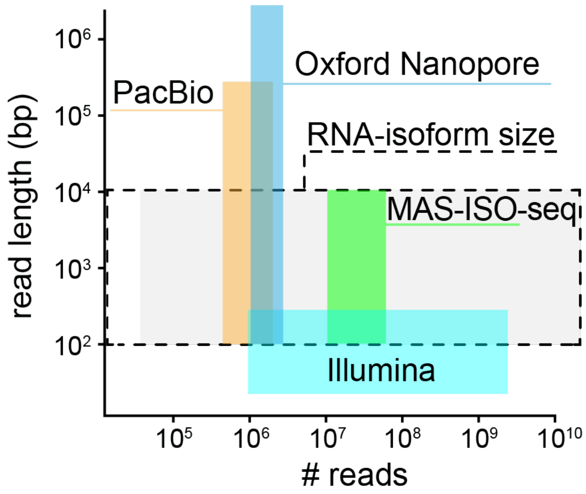


**Supplementary Fig 1. Read length and throughput of sequencing approaches.** Schematic displaying the read length and throughput of existing sequencing approaches and the context of RNA-isoform size.

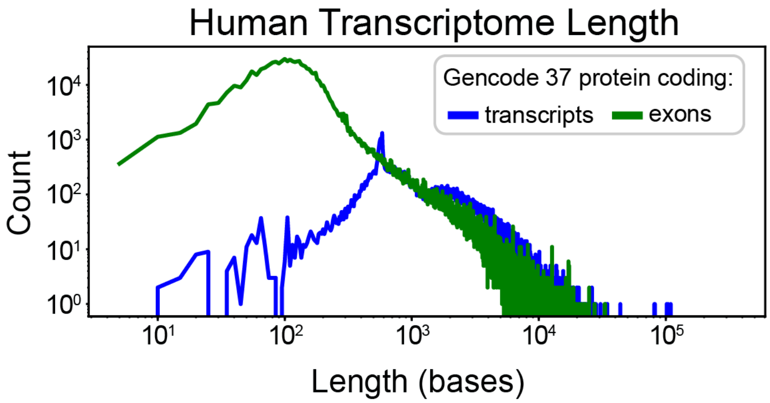


**Supplementary Fig 2. Human transcript/exon length distributions and statistics.** Histogram of human protein-coding transcript and exon length (obtained from Gencode version 37, HG38). The median protein-coding transcript length is 1595 (IQR: 2272.25) bases. The median protein-coding exon length is 127 (IQR: 98) bases.

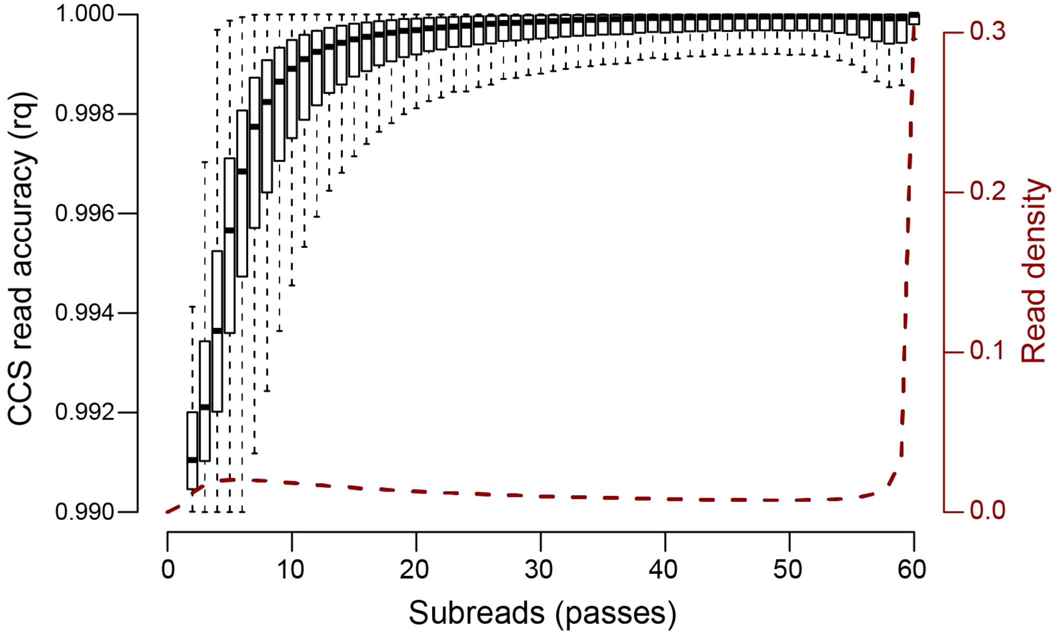


**Supplementary Fig 3. Error correction accuracy and circular pass frequency.** Left axis: software-estimated accuracy of circular consensus sequencing error correction as a function of number of circular passes. Right axis: fraction of overall reads with given number of circular consensus passes.

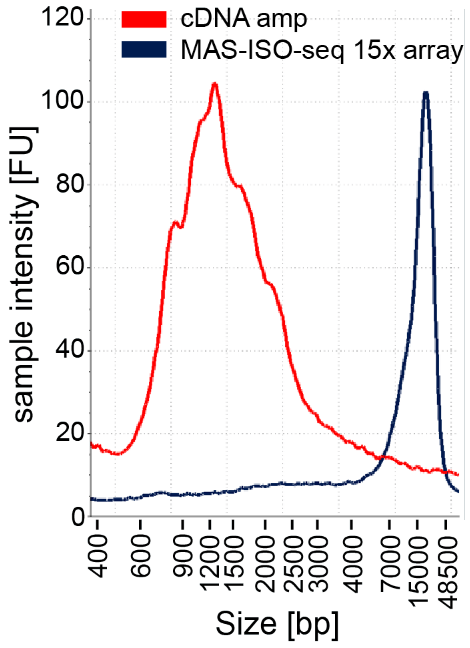


**Supplementary Fig 4. MAS-seq cDNA array size distribution.**
Electropherogram traces of cDNA library (red) and MAS-seq 15x cDNA array library (blue).

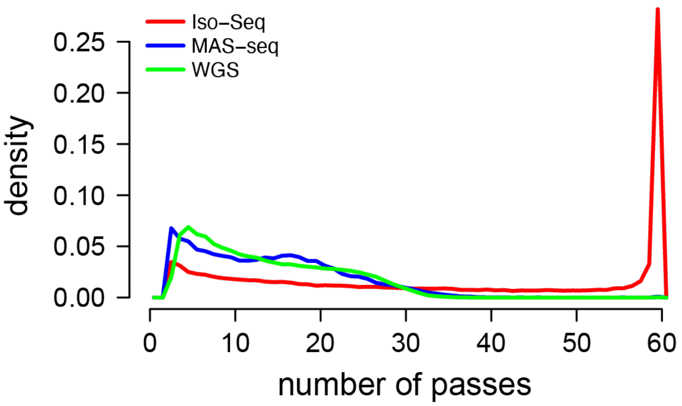


**Supplementary Fig 5. Comparison of subread circular consensus pass distributions across different PacBio sequencing modalities.** Red: SIRV set 4 isoforms sequenced via the Iso-Seq protocol. Blue: SIRV set 4 isoforms sequenced via the MAS-seq protocol. Green: PacBio WGS CCS data on HG002 (SRA accession SRR10382244).

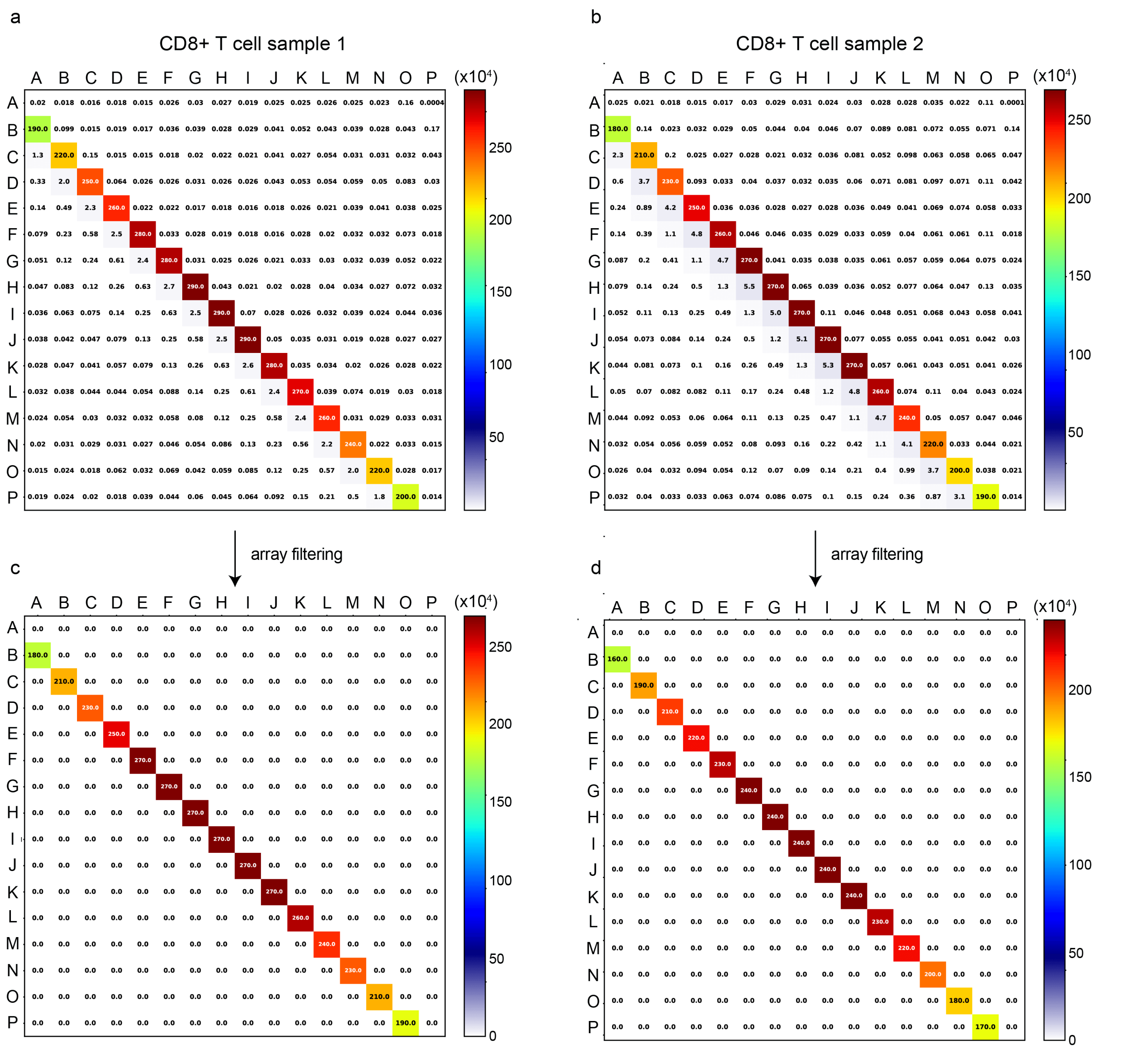


**Supplementary Fig 6. Ligation heatmaps for two T cell MAS-seq samples before and after filtering.** Heatmaps depict MAS-seq adapter pair frequencies as determined by Longbow annotations in two samples (left: M131TS, right: M132TS). Top row: before off-subdiagonal filtering. Bottom row: after off-subdiagonal filtering.

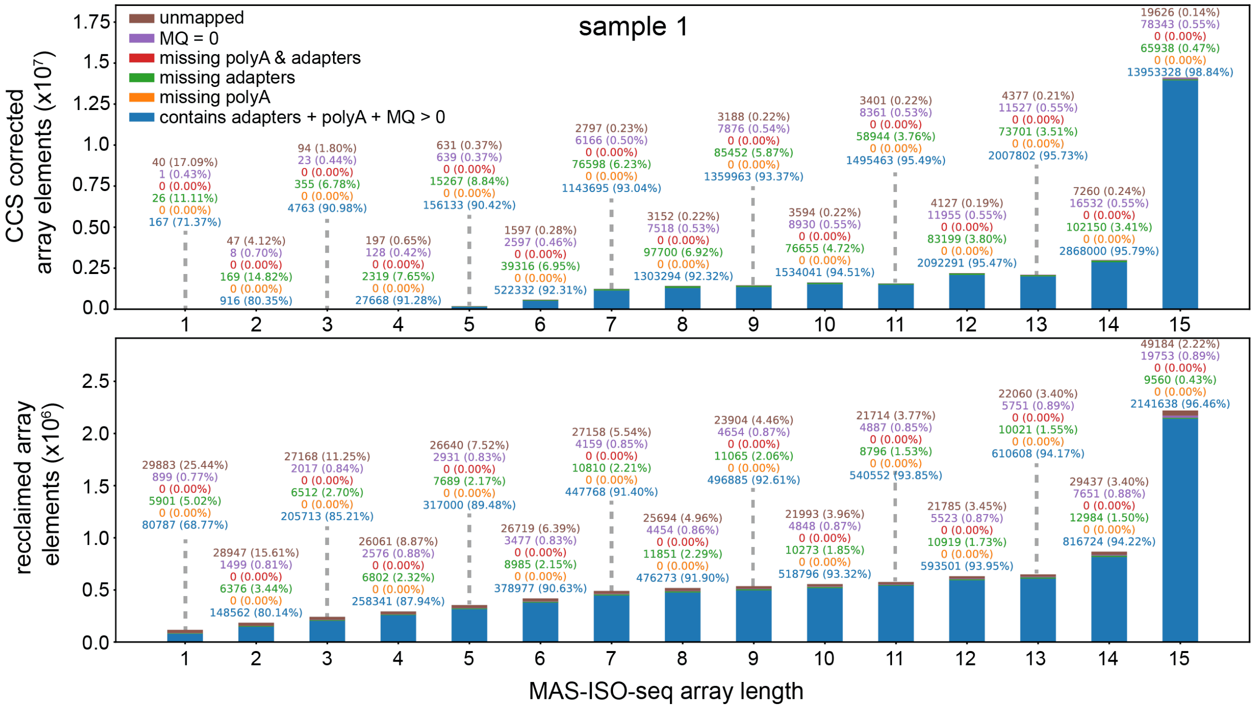


**Supplementary Figure 7. Segmented read classifications by MAS-seq array length (M131TS).** Counts of post-filtering/segmentation MAS-seq transcript reads originating from arrays of various lengths. **Top:** transcript reads that have been CCS corrected. **Bottom:** transcript reads that are uncorrected by CCS ("reclaimed"). Bar counts are broken into categories based on mapping statistics associated with the primary alignment of each read (blue: reads containing MAS-seq adapters and a poly(A) tail with mapping quality > 0, orange: reads missing a poly(A) tail, green: reads missing MAS-seq adapters, ed: reads missing MAS-seq adapters and poly(A) tails, purple: reads with mapping quality zero, brown: unmapped reads).

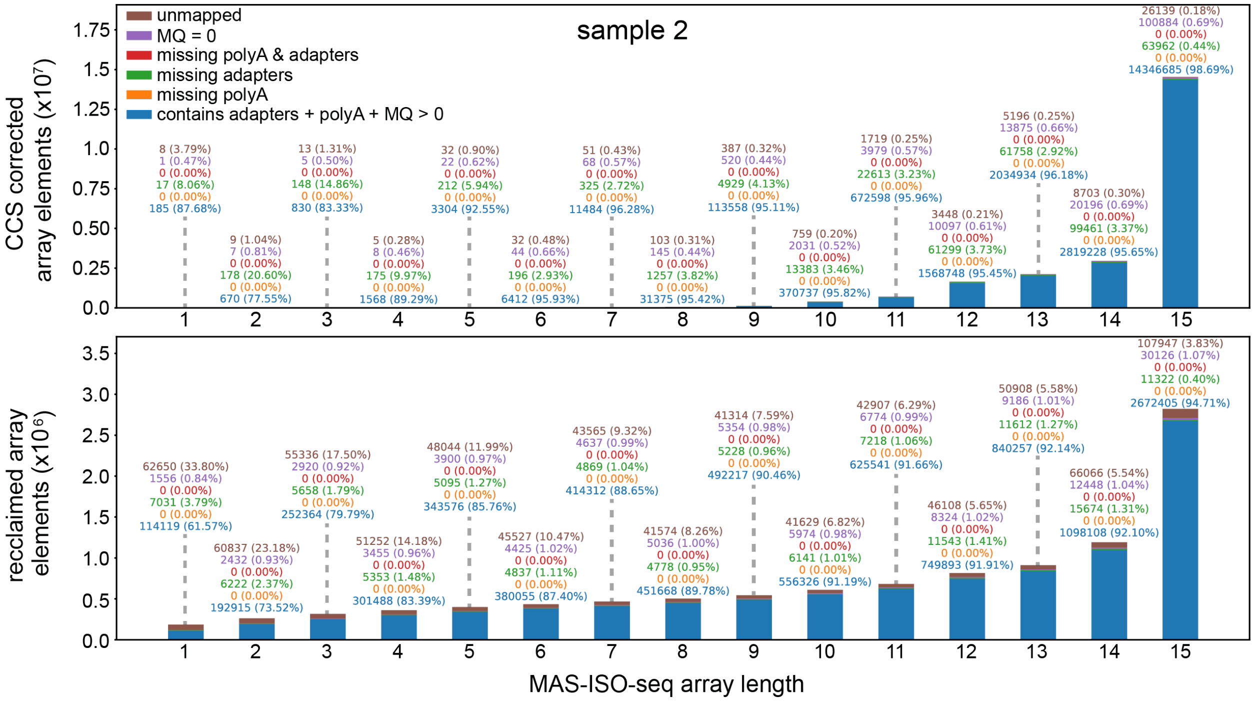


**Supplementary Figure 8. Segmented read classifications by MAS-seq array length (M132TS).** Counts of post-filtering/segmentation Longbow filtered MAS-seq array elements (transcript reads) originating from arrays of various lengths. **Top:** transcript reads that have been CCS corrected. **Bottom:** transcript reads that are uncorrected by CCS ("reclaimed"). Bar counts are broken into categories based on mapping statistics associated with the primary alignment of each read (blue: reads containing MAS-seq adapters and a poly(A) tail with mapping quality > 0, orange: reads missing a poly(A) tail, green: reads missing MAS-seq adapters, red: reads missing MAS-seq adapters and poly(A) tails, purple: reads with mapping quality zero, brown: unmapped reads).

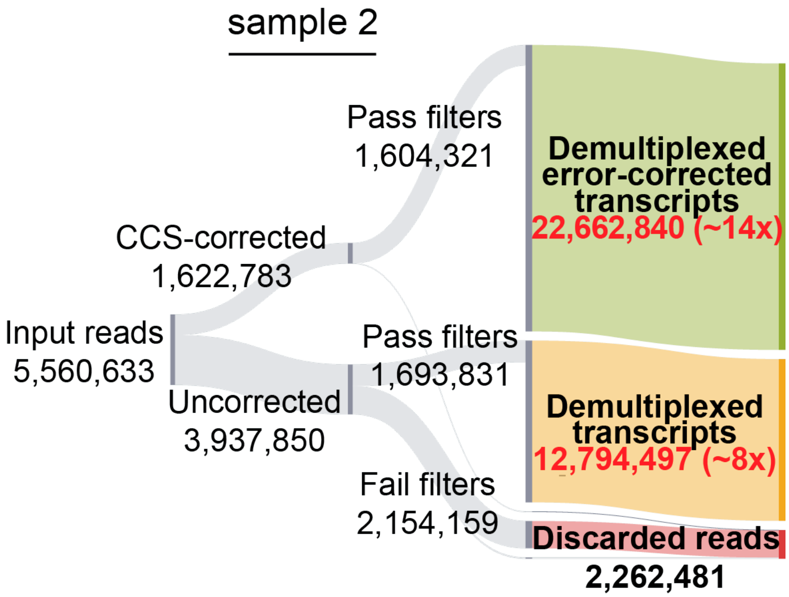


**Supplementary Figure 9. Sample 2 MAS-seq read metrics.** Sankey diagram reporting MAS-seq run yield of sample 2 at various stages of processing.

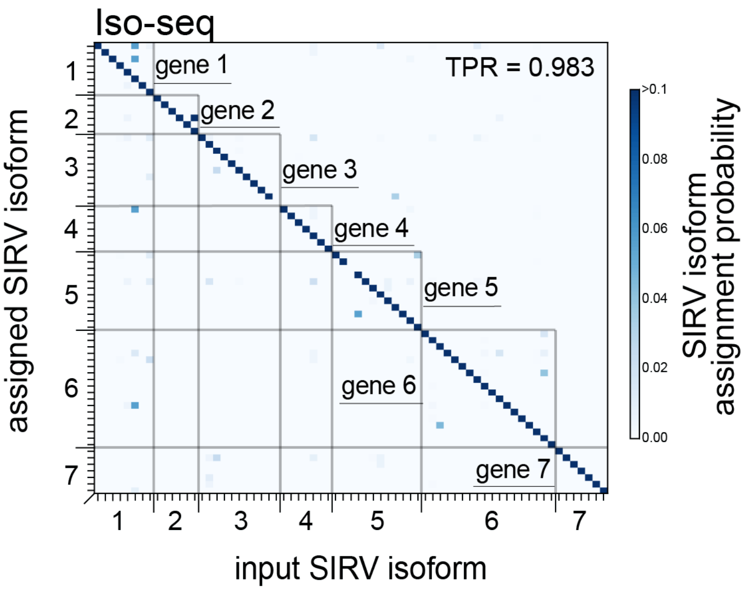


**Supplementary Figure 10. Synthetic RNA-isoform assignment - Iso-Seq.** Isoform identification confusion matrix for SIRV isoforms as measured by Iso-Seq.

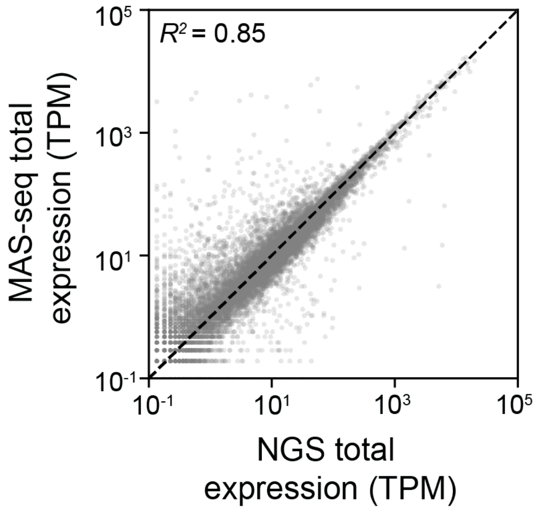


**Supplementary Figure 11. Gene expression concordance between MAS-seq long-read and Illumina short-read for CD8+ T cell dataset.** The total gene expression for MAS-seq is obtained by aggregating counts from all isoforms of every gene. The abundances are converted to transcript-per-million (TPM) counts for both MAS-seq and Illumina.

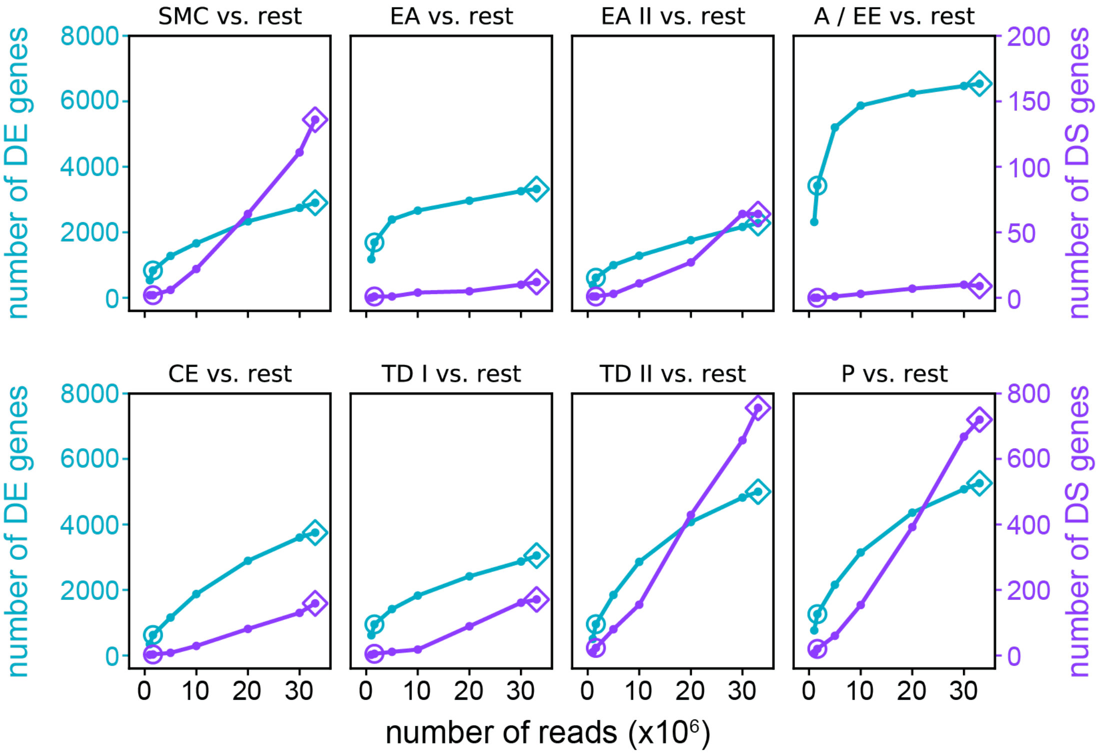


**Supplementary Figure 12. Downsampling analysis of MAS-seq CD8+ T cell dataset resolved by T cell subtypes.** The panels show the number of differentially expressed (DE) and differentially spliced (DS) genes for different T cell subtypes vs. read depth. The MAS-seq and Iso-Seq equivalent read depths are indicated by diamond (◇) and circle (○) makers. Note that identification of DE genes is nearly saturated for MAS-seq, and the increased read depth afforded by MAS-seq compared to Iso-Seq leads to vastly increased number of identified DS genes.

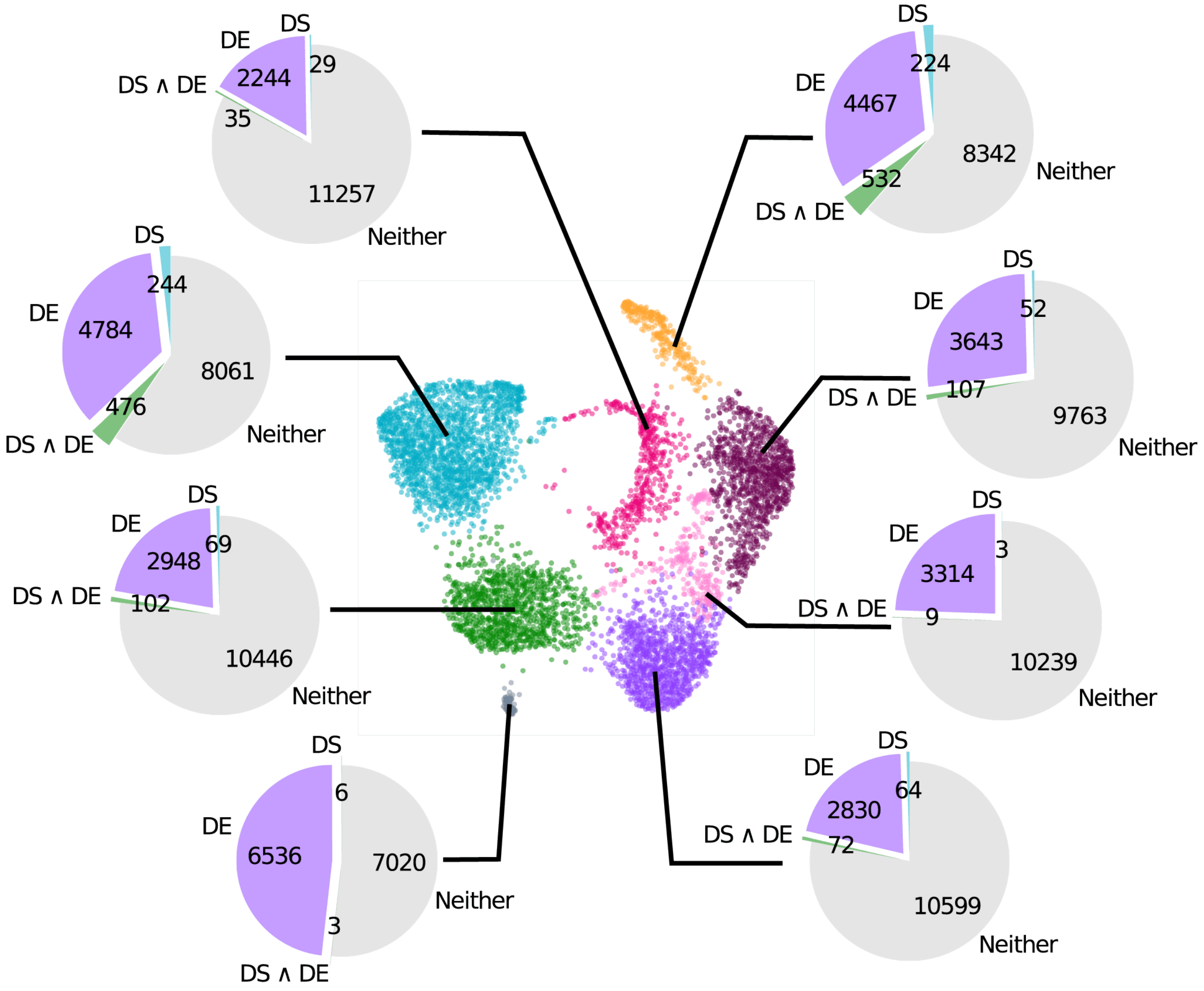


**Supplementary Figure 13. Number of differentially expressed (DE) and differentially spliced (DS) genes for MAS-seq CD8+ T cell dataset resolved by T cell subtypes.** We note that a plurality of identified DS genes are distinct from DE genes (as available to conventional short-reads scRNA-seq).

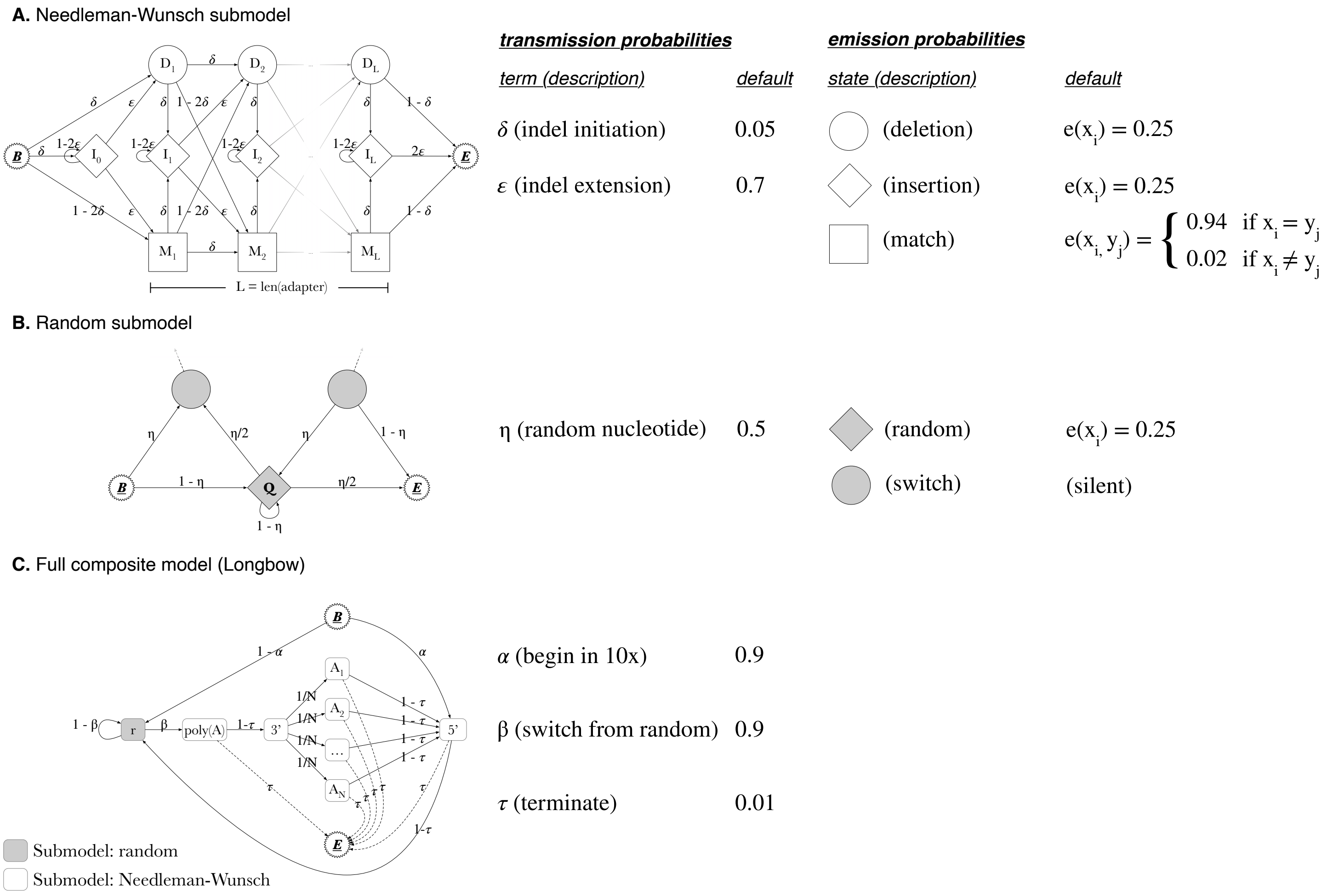


**Supplementary Fig. 14. Diagram of composite hidden Markov model, Longbow. a.** Needleman-Wunsch submodel for global alignment of a read subsequence to a known adapter of length L, allowing for mismatch, insertion, and deletion errors. Default indel initiation/extension transition probabilities and emission probabilities for match, insertion, and deletion states are annotated alongside the model diagram. **b.** Random sequence submodel for annotation of subsequences not known *a priori*. Transition probability into a random nucleotide, and emission probabilities for random and switch state (a silent state used to transition into other models) are annotated alongside the model diagram. **c.** Fully composed model for Longbow. White boxes indicate Needleman-Wunsch submodels. Grey box indicates random sequence submodel. Transition probabilities link model components in local order (i.e. 10x Genomics 5’ adapter, random (cDNA) sequence, poly(A) tail, 10x Genomics 3’ adapter, and one of N possible MAS-seq adapters where N is the number of array elements expected by the library design). Transitions to MAS-seq adapters are equally weighted (at this stage, MAS-seq ordering is not yet inspected or enforced) and permitted to transition to an end-state early, enabling the processing of partial arrays. Transition probabilities starting in the 10x Genomics 5’ adapter, switching from the random model, and termination are provided alongside the model diagram.

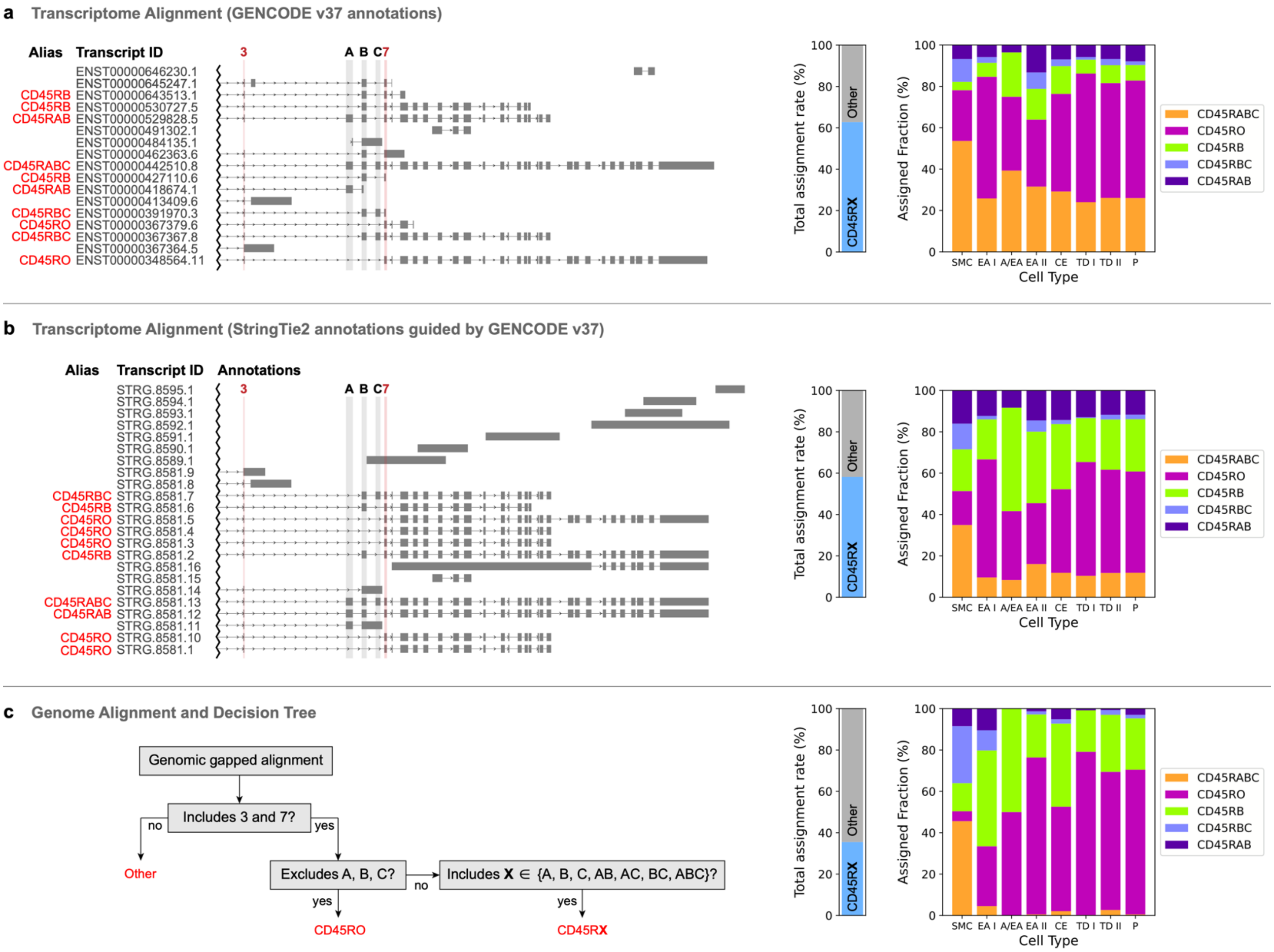


**Supplementary Figure 15. The effect of different isoform quantification strategies on CD45 isoform assignments to different CD8+ T cell subtypes. (a)** Alignment to GENCODE v37 transcriptome using minimap2 (splicing mode disabled). The annotations are shown on the left. Note that only ENST00000348564 (CD45RO) and ENST00000442510 (CD45RABC) extend to the 3' UTR whereas other annotations are incomplete. Consequently, most reads are primarily mapped to these two contigs regardless of the T cell subtype of origin (right). **(b)** Refinement of the GENCODE v37 annotations using StringTie2 based on MAS-seq reads, followed by alignment to the refined transcriptome using minimap2 (splicing mode disabled). Note the increased completeness of the annotations and subsequently (left), the increased specificity of isoform assignments (right). **(c)** Genome alignment of MAS-seq reads to GRCh38 using minimap2 (splicing mode enabled) followed by isoform assignment using a manually defined decision tree (left). Note the further increased specificity of isoform assignments.

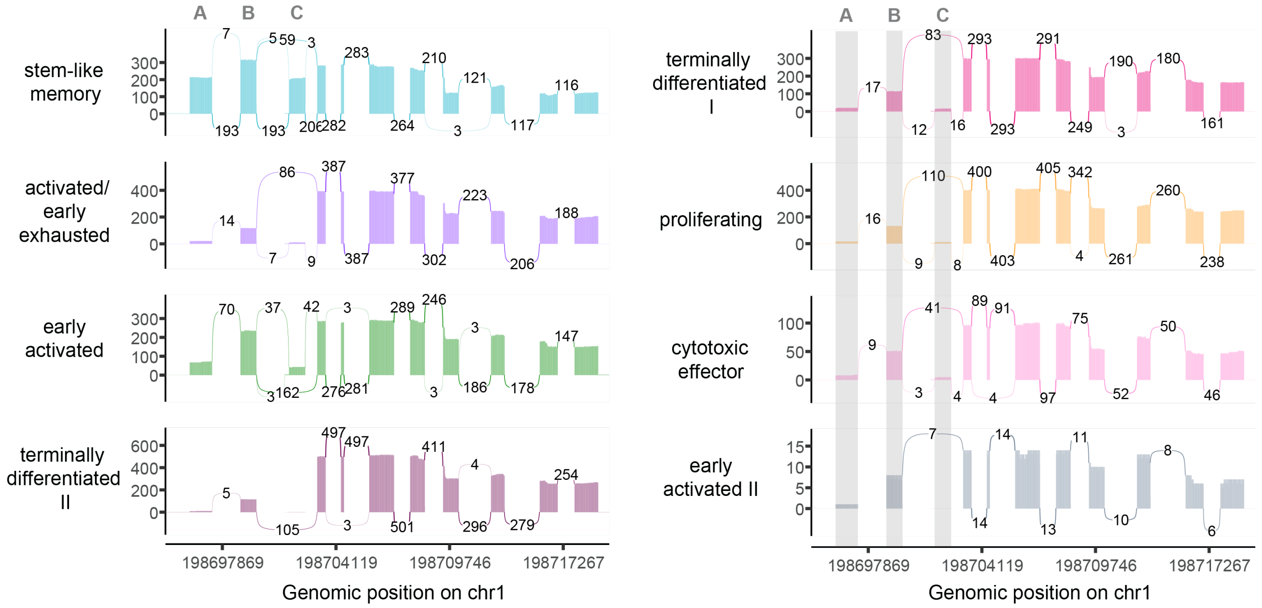


**Supplementary Figure 16. Sashimi plot of CD45 full-length transcripts in MAS-seq CD8+ T cell dataset resolved by T cell subtypes.** The plot is produced using ggsashimi package(Garrido-Martín et al. 2018). Note the highly variable inclusion/exclusion pattern of the landmark exons A, B, and C. Notably, the CD45RABC isoform (including all three exons) is prevalent in stem-like memory cells whereas the CD45RO isoform (excluding all three exons) is prevalent in activated and terminally differentiated subtypes.
